# Immunosuppression broadens evolutionary pathways to treatment failure during *Acinetobacter baumannii* pneumonia

**DOI:** 10.1101/2021.04.07.438861

**Authors:** Wenwen Huo, Lindsay M. Busch, Juan Hernandez-Bird, Efrat Hamami, Christopher W. Marshall, Edward Geisinger, Vaughn S. Cooper, Tim van Opijnen, Jason W. Rosch, Ralph R. Isberg

## Abstract

*Acinetobacter baumannii* is increasingly refractory to antibiotic treatment in healthcare settings. As is true of most human pathogens, the genetic path to antimicrobial resistance (AMR) and the role that the immune system plays in modulating AMR during disease are poorly understood. Here we reproduced several routes to fluoroquinolone resistance, performing evolution experiments using sequential lung infections in mice that are replete or depleted of neutrophils, providing two key insights into the evolution of drug resistance. First, neutropenic hosts acted as reservoirs for the accumulation of drug resistance during drug treatment. Selection for variants with altered drug sensitivity profiles arose readily in the absence of neutrophils, while immunocompetent animals restricted the appearance of these variants. Secondly, antibiotic treatment failure in the immunocompromised host was shown to occur without clinically defined resistance, an unexpected result that provides a model for how antibiotic failure occurs clinically in the absence of AMR. The genetic mechanism underlying both these results is initiated by mutations activating the drug egress pump regulator AdeL, which drives persistence in the presence of antibiotic. Therefore, antibiotic persistence mutations present a two-pronged risk during disease, causing drug treatment failure in the immunocompromised host while simultaneously increasing the emergence of high-level AMR.

## Introduction

*Acinetobacter baumannii* is a Gram-negative opportunistic pathogen, one of the high-priority ESKAPE organisms that are increasingly difficult to treat due to multiple antibiotic resistance^1^. A significant proportion of healthcare-associated diseases caused by this group of pathogens, such as ventilator associated pneumonia, is linked to their documented multi-drug resistance (MDR) ^2–5^. Of particular importance are patients in intensive care units (ICU) who are critically ill and have depressed immunological clearance mechanisms that increase the risk of infection by MDR pathogens ^6, 7^. As a consequence, the healthcare environment with its immunologically compromised patients could provide a unique niche for selection of MDR isolates ^8^. Overlaying these issues is the fact that for many patients in healthcare settings, antibiotic treatment failure is common but is often unexplained, as resistant organisms cannot be identified^9^.

*A. baumannii* clinical isolates have demonstrated a remarkable ability to successfully battle antibiotic treatment in the clinic, resulting from high intrinsic resistance to antimicrobials and the acquisition of drug resistance elements by the organism^10, 11, 12, 13^. A critical missing link is a detailed roadmap for the stepwise evolution of antibiotic resistance in the clinic, particularly in identifying *A. baumannii* subpopulations most likely to give rise to drug treatment failure. Furthermore, it is largely unknown if there exists a patient group that provides the reservoir for antimicrobial resistance (AMR) acquisition. Particularly for healthcare-associated diseases, patient groups susceptible to *A. baumannii* are by their nature compromised in a number of fashions, with the potential for providing reservoirs for AMR evolution. The range of individuals with altered immune function in these settings may allow for a diversity of host targets that can act as primary amplifiers of resistance, with eventual spread to individuals with different sets of susceptibilities. Therefore, as a model for healthcare-associated pneumonia we aimed to explore whether depletion of a single arm of innate immunity in mice could help shape the antibiotic treatment outcome and support the evolution of resistant organisms.

Fluoroquinolones (FQ) such as ciprofloxacin initially showed excellent activity against *A. baumannii* infections^14^. Members of this drug class inhibit bacterial cell growth by covalently linking to DNA gyrase (*gyrA*) and topoisomerase IV (*parC*), leading to double stranded DNA breaks and cell death^15^. Drug resistant mutants arise fairly frequently in the clinic, with over 80% of clinical isolates of *A. baumannii* currently being FQ resistant^16^. The most commonly reported mechanisms of ciprofloxacin resistance in the clinic are target protein alterations and the overexpression of efflux pumps. In *A. baumannii*, alterations in target proteins usually evolve in a stepwise fashion, starting with *gyrA* (usually Ser81Leu) followed by *parC* (Ser83Leu). Interestingly, in addition to the patient’s underlying condition and hospitalization status, prior exposure to fluoroquinolones is also a risk factor for *A. baumannii* colonization and infection^17^, indicating that resistance to this antibiotic class is linked to either increased pathogenic potential of the isolate or is highly associated with acquisition of MDR. Unknown is whether there exist early adaptive mutations that enable precursor populations of *A. baumannii* to act as ancestors to drug resistance.

There has been little study of whether AMR can be suppressed by the immune response. Landmark mathematical modeling work argues that the immune response can largely limit the outgrowth of persisters or other bacterial variants that exhibit intermediate resistance levels ^18^. Another study indicates that the cytokine response may control waves of AMR variants^19^. Given the limited analysis of how antibiotic resistance evolves in the clinic, and the lack of a detailed interrogation of the role played by the immune response in controlling selection of AMR, we sought to identify the steps that lead to resistance in the presence or absence of a single arm of innate immunity. The rationale behind this approach is that clinical antibiotic resistance is associated with mutations located outside well-characterized drug targets, and these are difficult to identify bioinformatically or predict based on culture studies ^20–22^. This approach provides evidence that absence of neutrophil function allows outgrowth of drug persisters and the appearance of fluoroquinolone resistance. In so doing, we identified mutational pathways to drug resistance, with the results tied to the problem of unexplained antibiotic treatment failure.

## Results

### Recapitulation of *Acinetobacter baumannii* evolved drug resistance during pneumonic disease

We serially passaged *A. baumannii* 15 times within a mouse pneumonia model to analyze the dynamics and genetic trajectories of resistance to the FQ antibiotic ciprofloxacin (CIP), with the purpose of determining whether neutrophils play a role in suppressing drug resistance (Fig. 1A). The CIP^S^ ATCC reference strain 17978 (AB17978) was passaged by oropharyngeal inoculation in either immunocompetent animals or those depleted of neutrophils by pretreatment with two doses of cyclophosphamide. It should be noted that this treatment also has other effects on the immune system, including the possible reduction of monocyte populations, as well as suppressing T-cell numbers ^23, 24^. Mice in three parallel lineages were CIP treated at 7- and 19- hours post infection (hpi) and bacteria were collected from the lungs of animals euthanized at 27 hpi. The enriched bacterial pools from each independent infection were used as inocula for the next round of infection. The dynamics of bacterial yield was assessed from lung homogenates, and the CIP minimum inhibitory concentration (MIC) of each population was determined followed by whole genome sequencing of the heterogenous pool (mean genome coverage depth: 368.4).

**Figure 1.**
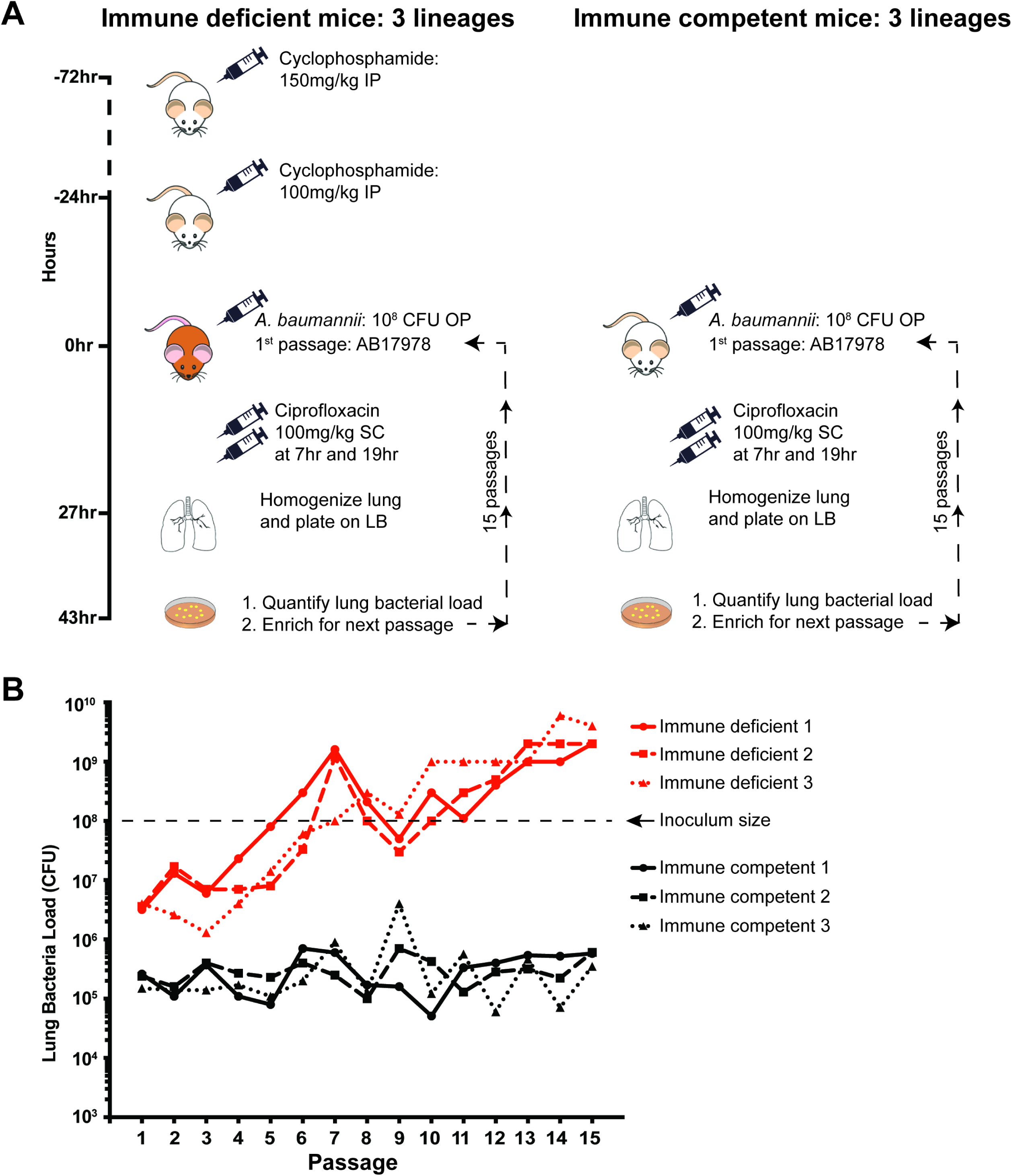
Continuous passaging of *A. baumannii* in a lung infection model results in early antibiotic failure in neutrophil-depleted mice. A). The 15-passage strategy using the pneumonia model in both immunocompetent and neutrophil-depleted mice. Neutrophil depletion was induced by two doses of cyclophosphamide at 72- and 24-hrs prior of infection. At the time of infection, *A. baumannii* ATCC 17978 was oropharyngeally inoculated into mice at 10^8^ CFU, and two doses of CIP were administered at 9- and 19-hrs post infection. All Lineages are derived from a single broth-grown culture. At 27-hrs post infection, the mice were euthanized, lungs were homogenized, and bacteria were plated on LB agar plates. After overnight growth, the bacteria were collected and 10^8^ CFU used for the next round of infection/passage. B). Rapid development of antibiotic failure as a consequence of passage in the absence of neutrophils. After lungs were homogenized, total CFU on LB agar in the absence of antibiotic were determined and plotted as function of passage number. Shown are three lineages for immunocompetent and neutrophil-depleted animals treated with CIP as described in panel (A).

### Persistent neutropenia during successive passages quickly results in antibiotic treatment failure

A striking result from the passage experiments is that the protocol allowed drug resistant mutants to overgrow during passage in neutrophil-depleted animals, while the resistant variants in the immunocompetent animal failed to reach high abundance in the population. After each mouse passage in the presence of CIP, the efficiency of bacterial colonization of the lungs was monitored by quantifying total bacterial colony forming units in the absence of drug (CFU). Bacterial lung yields increased by 1000-fold in CIP-treated neutrophil-depleted mice over the course of 15 passages, as compared to little increase in immunocompetent mice (Fig. 1B). After each passage, we observed 10-fold higher colonization in the neutrophil-depleted compared to immunocompetent mice. By passages 5-7, the lung bacterial load in the neutrophil-depleted mice reached the size of the initial inoculum (10^8^ CFU), and animals showed significant signs of disease such as lacrimation, piloerection, and decreased mobility in spite of being CIP treated (Fig. 1B).

To determine levels of CIP resistance after each passage, the saved bacteria were quantified on solid medium in the absence or presence of increasing amounts of drug, plotting the fraction of surviving CFU as a function of CIP concentration (Material and Methods; Fig. 2A and 2C). Clinically resistant bacteria (CIP ≥2ug/ml) did not arise in the three Lineages passaged in immunocompetent mice. Furthermore, the number of bacteria able to survive on increased amounts of CIP relative to the parental strain never rose above 0.1% of the population (survival on solid medium containing 1ug/ml CIP). Therefore, in immunocompetent mice, 15 passages were insufficient to allow outgrowth and fixation of resistant variants in the population.

**Figure 2.**
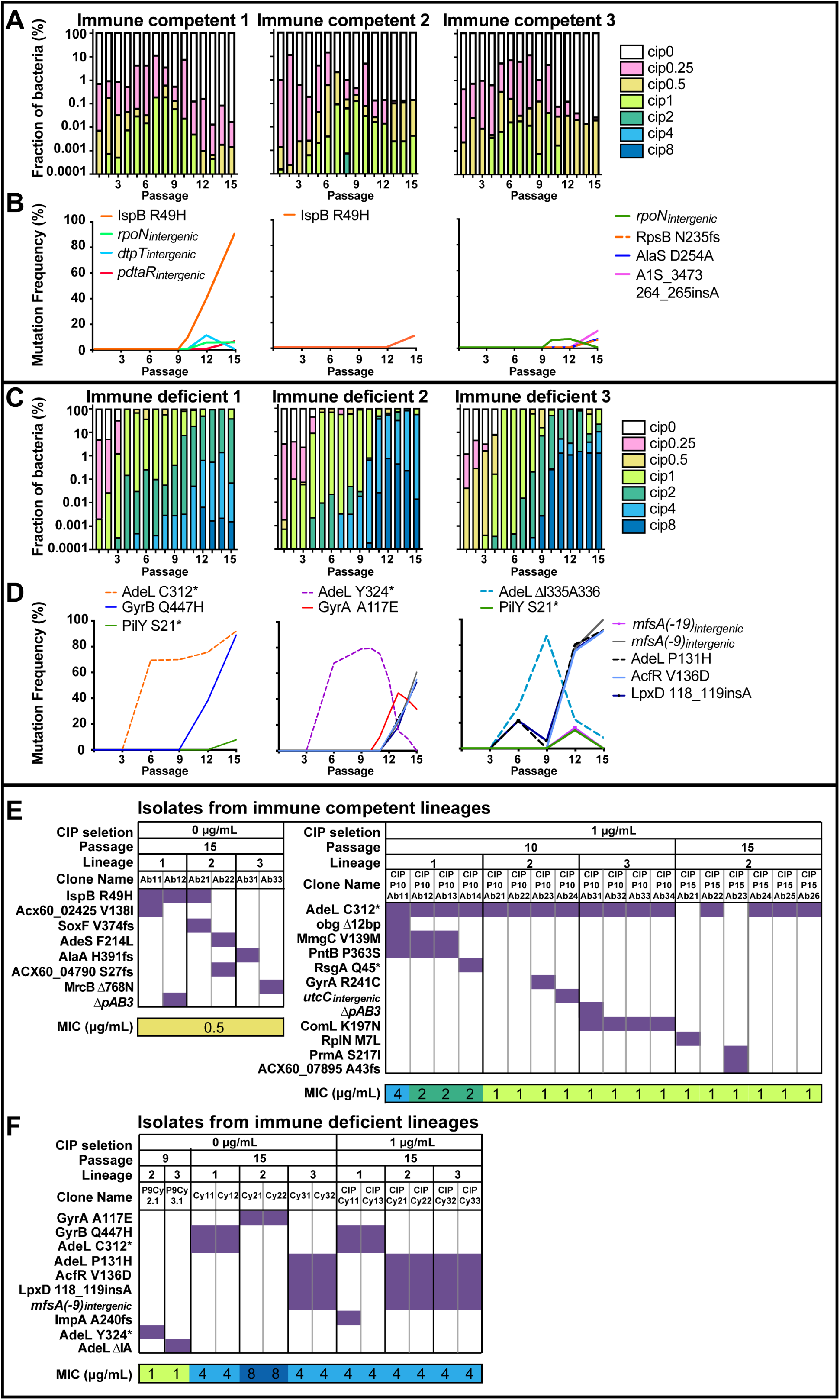
High level ciprofloxacin resistance evolves in a stepwise fashion during passage in neutrophil-depleted mice. (A, C) After each passage, bacteria from lung homogenates were serially diluted onto LB agar containing noted concentrations of CIP (μg/ml). The fraction of CFU on the series of CIP plates was determined for each pool and displayed in stacked bar plots for immune competent (A) and immune depleted (C) mouse infection Lineages. The limit of detection was 2.7E-6. (B, D) The gDNA from each bacterial pool was isolated and subject to whole genome sequencing to identify the genomic mutations acquired throughout passaging (after passage 3, 6, 9, 12 and 15). The mutations were detected using a 5% abundance cutoff and filtered against the parent strain AB17978. The relative abundance of each mutation was plotted as a function of passage number from (B) immune competent Lineages and (D) immune deficient Lineages (D). (E-F) After 10 or 15 passages (as noted), the bacterial pools from each Lineage were incubated on LB in the presence or absence of CIP (1 μg/mL) to select for single colonies. Purified single colonies were subjected to whole genome sequencing and MIC assays. In (E-F), each column represents a single colony. Rows display absence or presence of selection criteria, passage, amino acid or SNP alleles, and MIC values for each single colony isolate. The presence of each mutation in an individual colony is highlighted in purple. MICs are labeled and color-coded for all colonies, a minimal of three biological replicates were performed.

The evolutionary trajectories through neutrophil-depleted mice contrasted strongly with immunocompetent mice, as the whole population in all three Lineages became CIP resistant over the course of 15 passages. Bacteria resistant to at least 2 μg/ml CIP were observed as early as the 3^rd^ passage (Fig. 2C). After the 9^th^ passage, approximately 1x10^-5^ of the bacteria isolated from the neutrophil-depleted mice showed resistance to 8 μg/mL CIP, four times higher than the clinically-defined resistance breakpoint. By 15 passages, almost all bacteria from the neutrophil-depleted Lineages were clinically defined as CIP resistant (at concentration of at least 2 μg/mL), with a large fraction showing resistance to CIP =8 μg/ml. Therefore, resistance acquisition during disease is amplified in the absence of neutrophils.

### Clinical failure results from a bacterial population that is clinically defined as drug sensitive

By the 6^th^ passage in neutrophil-depleted mice, yields of bacteria began to approach those observed in the absence of antibiotic (Fig. 1B), in spite of the fact that the majority of the population remained sensitive to CIP based on the defined clinical breakpoint (Fig. 2C; clinical breakpoint CIP>2 μg/ml). Yields remained high in successive passages, while the distribution of the population increasingly skewed toward variants with increasing levels of resistance to CIP. Therefore, drug treatment failure at passage 6 under neutrophil-depleted conditions was dominated by bacteria that grew at 1μg/ml CIP but showed low viability at the 2μg/ml clinical breakpoint. These mutants were replaced by more resistant isolates during later passages (Fig. 2C).

To link specific mutational changes over time to these observed phenotypes, we sequenced the pools of genomic DNA isolated from the original inoculum, the parent AB17978 strain, and the pools harvested after the 3rd, 6th, 9th, 12th and 15th passages. Mutation frequencies were then plotted for each passage in the two immune conditions (Fig. 2B and 2D). Based on the predicted sensitivity from the depth of pooled whole genome sequencing, we expected that only nucleotide changes in greater than 5% of the population could be detected using this approach (Materials and Methods).

The results from the pooled genomic sequencing allowed us to identify changes associated with either the development of drug resistance, or the lack thereof, in these populations. As predicted from Fig. 2A and the sensitivity limits of the analysis, there was no evidence for mutations driving drug resistance at greater than 5% abundance in the three populations evolved in immunocompetent mice. The only alteration that approached fixation was a mutation in a gene encoding a putative solanesyl diphosphate synthase (ACX60_03850; *ispB*), but this allele was unrelated to resistance, as single colony isolates showed no increased survival in the presence of CIP (Fig. 2E). All the other mutations identified by sequencing the pools appeared unrelated to resistance, as none survived incubation on solid medium containing 1μg/ml CIP (Figs. 2B, 2E). Furthermore, the analysis of individual colonies isolated after plating bacterial populations on antibiotic-free medium after the 15^th^ mouse passage identified isolates with CIP MICs that were only slightly elevated relative to the WT (Fig. 2E, left panel; WT MIC = 0.25-0.5).

Variants with altered CIP sensitivity arose in the immunocompetent mice, but they remained at low levels in these populations, as predicted by the previous mathematical modeling study (Fig. 2A) ^18^. Mutations causing decreased drug sensitivity were identified by isolating single colonies on solid medium containing 1μg/ml CIP, and all twelve of the single colonies selected from the 10^th^ passage had the identical mutation in *adeL*, the regulator of the AdeFGH egress pump ^25, 26^. The presence of the mutation in multiple Lineages is consistent with it existing in the initial bacterial culture prior to the first inoculation of all three Lineages. Interestingly, 8 distinguishable Lineage-specific strains were identified that were derived from the parental AdeL C312* mutation, consistent with downstream mutations arising during mouse passage (Fig. 2E, right panel). These strains appeared to be largely lost by passage 15 in the immunocompetent mouse (Fig. 2A, E). We conclude that although strains with mutations that activate the AdeFGH egress pump are detected, they are unable to overgrow the WT and are eventually depleted during later passages in immunocompetent mice (Fig. 2B).

Consistent with the dynamic nature of resistance acquisition in neutrophil-depleted mice (Fig. 2B), pool sequencing uncovered mutations that drove the stepwise trajectory to CIP^R^ in later passages (Fig. 2D). Unlike the immunocompetent lineages, first step mutations that occurred within *adeL* after 6 passages in neutrophil-depleted mice were easily detected by whole pool sequencing. Each Lineage harbored different *adeL* mutations in these early passages that disrupted the 3’ end of the gene, consistent with previous work arguing that alterations in the C-terminus of the AdeL protein result in AdeFGH pump activation^25, 26^. Mutations in various other genomic sites arose between passage 9 and 12, with many predicted to contribute to decrease susceptibility to CIP treatment. For instance, the *adeL312\** mutant acquired a second mutation in *gyrB* that overgrew Lineage 1. In addition, a mutant harboring a noncononical mutation in *gyrA* arose in Lineage 2 that was outcompeted by a strain with multiple genetic alterations, while a quadruple mutant appeared to overgrow the single *adeL* mutant in Lineage 3.

To identify the various genotypes linked to increased CIP resistance, we isolated single colonies after 9 passages to verify that two transiently predominant variants had the predicted genotypes, and also sequenced isolates from passage 15 after plating in the absence or presence of CIP (1µg/mL). Isolated colonies in absence of drug selection from Lineages 2 and 3 from passage 9 showed the predicted *adeL* alleles (compare Fig. 2D and 2F). In addition, after plating the passage 15 pools in the absence of drug, the two predicted gyrase alleles *gyrA*(A117E) and *gyrB*(Q447H) that are observed infrequently in the clinic were also identified (Fig. 2F). Having *gyrA*(A117E) alone was able to raise the CIP MIC by 16-fold (Fig 2F). Clones having the *gyrB*(Q447H) allele were only observed linked to *adeL*(C312*) at passage 15, resulting in a CIP MIC that was 8-fold higher than that of the parent strain (Fig 2E). Besides the mutations in target proteins, other single colonies that resulted in a similar MIC increase were found to have a combination of four mutations (Fig. 2F). When comparing the single colonies isolated that had a CIP of 1 μg/mL to those isolated without antibiotic, the single colony isolates had genomic changes that were predicted by the sequencing of the pools, indicating that the mutants described here became predominant without requiring CIP selection *ex vivo*.

### Mutations in *adeL* allow for persistence in the presence of high levels of ciprofloxacin

To deconvolve the function of the various mutations observed, we backcrossed mutations into the parent strain and assessed their relative contributions to CIP resistance. Of particular interest were *adeL* nucleotide changes identified in populations after the 6^th^ passage in neutrophil-depleted mouse lineages, as CIP failed to efficiently restrict growth in the lung in these passages (Figs. 1A, 2D). The *adeL* gene (ACX60_06025)^26^ has two predicted domains often associated with LysR type transcriptional regulators: a helix-turn-helix domain and a substrate binding domain responsible for regulation of the AdeFGH pump (Fig. 3A). Of the *adeL* mutations that arose during the *in vivo* passages, one is located within the predicted substrate binding domain while the others are at the C terminal end comprising in-frame deletions or early termination codons (Fig. 3A and 3B). We constructed strains with each of these mutations and observed 100-1000X increased *adeG* transcription levels compared to the parental strain, consistent with AdeFGH efflux pump overexpression (Fig. 3C).

**Figure 3.**
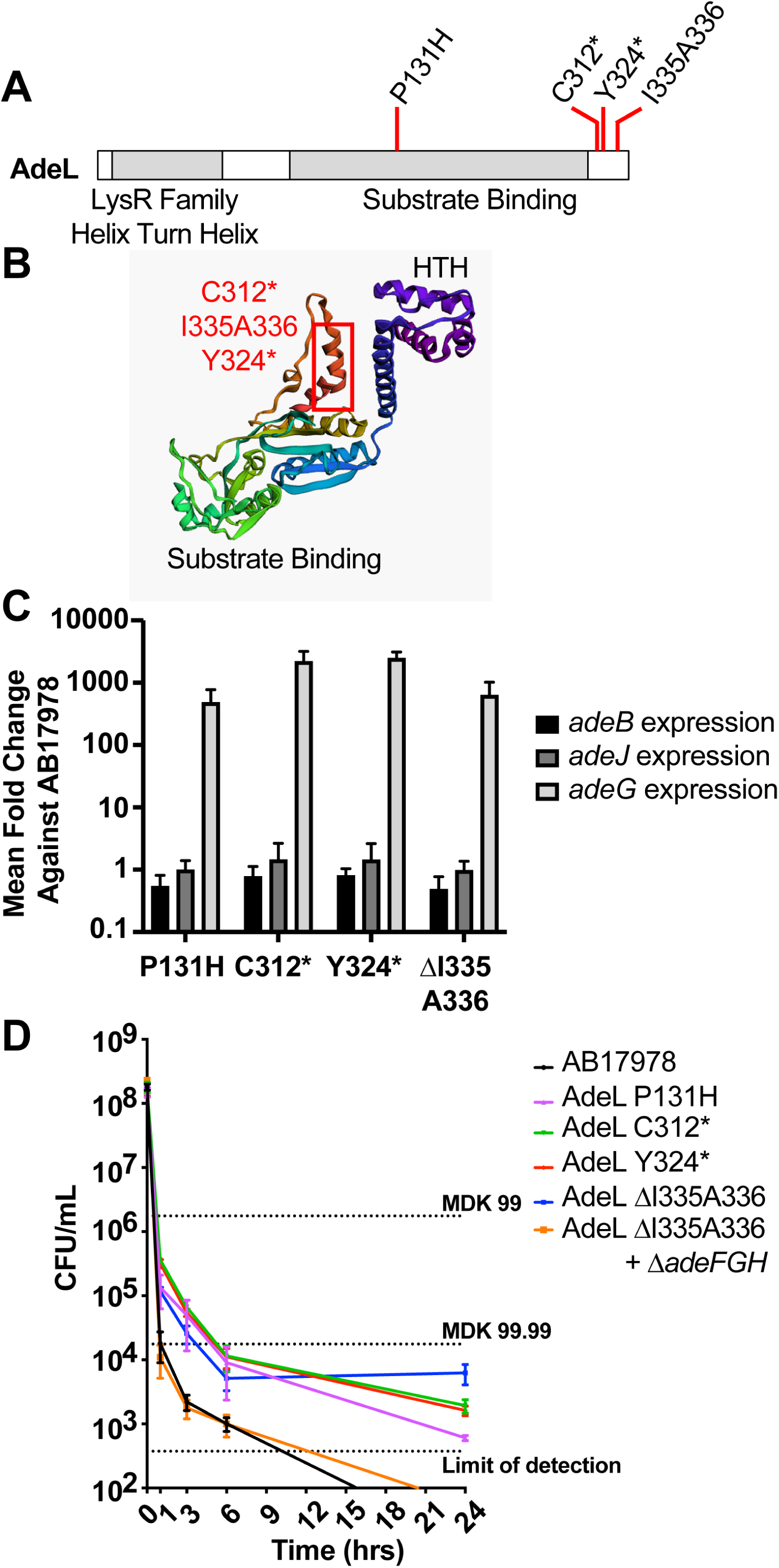
*adeL* mutations drive bacterial persistence through over-expression of the AdeFGH pump. (A). Sites of nonsynonymous changes during *in vivo* passaging and domain structure of AdeL. (B). Locations of nonsynonymous changes on predicted functional domains of AdeL. (C). Mutations in *adeL* selected during mouse passage result in specific overproduction of an AdeL-regulated gene *adeG*. Data were determined by qRT-PCR analysis of pump-encoding components *adeG* (regulated by AdeL) or *adeB* and *adeJ* (components of other pumps) in noted *adeL* mutant backgrounds. (D). Overproduction of AdeL-regulated pump components results in increased persistence in presence of CIP. AB17978 derivatives described in legend were exposed to CIP at 20X MIC for noted times and titered for viability by quantitating CFUs. Data shown are mean ± Standard Error of Mean, a minimal of three biological replicates were performed.

While these single *adeL* mutations resulted in an increase in MIC to ∼1 μg/mL CIP that barely rose to the level of significance (Fig. 4C), this level was below the recognized clinical breakpoint^27^, and such strains would be indicated as susceptible. As these are first step mutations, we hypothesize that variants harboring these changes may facilitate the outgrowth of more resistant isolates by promoting tolerance or persistence in the presence of antibiotic. To test this model, we evaluated the survival of *adeL* mutants during exposure to high levels of CIP ^28–32^. Exposure to 10 μg/mL CIP, roughly 20X MIC of the parental strain, led to the majority of the *adeL* mutants (> 99%; MDK99) being rapidly killed (within 1 hour), mimicking what was observed for the parental strain (Fig. 3D). However, unlike the parental strain, a subpopulation of the *adeL* mutant was able to persist through 24 hours of drug exposure (Fig. 3D). This phenotype was dependent on pump overproduction, as an *adeL* mutant strain deleted for the *adeFGH* operon was indistinguishable from the parental strain in regards to persistence (Fig. 3D). These data argue that *adeL* mutations drive persistence in the presence of CIP *via* increased expression of *adeFGH*, providing a reservoir within neutrophil-depleted animals for the outgrowth of strains with increased resistance^33^.

**Figure 4.**
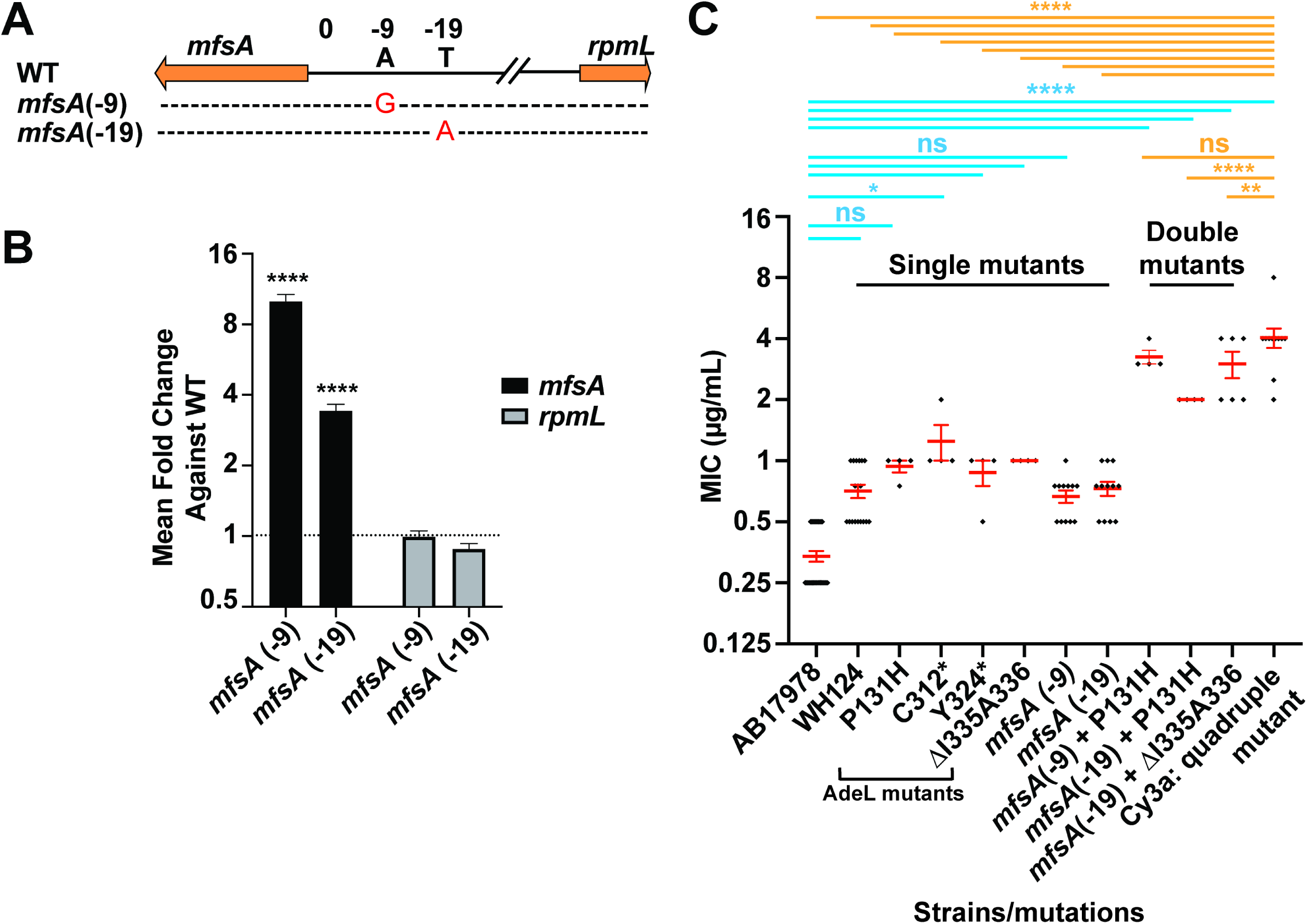
The presence of mutations in *adeL* and *mfsA* is sufficient to explain CIP^R^ in strains derived from mouse passage experiments. A) Two mutations located directly upstream of *mfsA* selected during mouse passaging experiments. B) Transcription levels of *mfsA* and *rpmL* were determined by qRT-PCR in strains harboring the two mutations upstream of *mfsA*. Significance level: ****, p < 0.0001 using two-tailed t-test against mean fold change of WT. Two technical replicates and at least three biological replicates were performed. Data are mean MICs ± SEM. C) Double mutants containing *adeL* and *mfsA* mutations show MICs of CIP that are similar to evolved quadruple mutant strain. Data are raw MIC values for each biological replicate in black with mean MICs ± SEM in red. At least three biological replicates were performed. WH124: AB17978 with *lpxD* insA *acfR* V136D *adeL* P131H ; C312*: Cysteine to stop codon at residue 312; Y324*: Tyrosine to stop codon at residue 324. Statistical significance was tested using one-way ANOVA followed by Dunnett’s multiple comparison. Blue: multiple comparison against WT AB17978 MICs; Orange: multiple comparison against MICs of Cy3a quadruple mutant. *: p<0.05; **: p<0.01; ****: p<0.0001; ns: not significant.

### Resistance of evolved mutants in neutrophil-depleted mice can be explained by combining *adeL* with *mfsA* mutations

We continued backcross analysis to identify the minimal determinants necessary to confer clinical resistance during passage in the mouse. Many of the single colony isolates from Lineage 3 at passage 15 of the immunocompromised mice possessed a combination of mutations in four genomic locations: *adeL*, *acfR* (ACX60_03155), *lpxD*, and the intergenic *rpmL*-*mfsA* region, the latter of which is located between a ribosomal protein gene and a predicted coding region for a putative MFS transporter which we have named *mfsA* (ACX60_15150, annotated as *nreB* in GenBank; Fig. 2D and 2E). To determine the role in drug susceptibility of each of these mutations, we constructed separate strains carrying single mutations. All single mutants showed various MICs well below 2 μg/ml CIP, with each of the *adeL* mutations as well as *mfsA*(-9) showing increased MICs above the parental strain (Table S1; Fig. 4C). The *mfsA*(-9) mutation is located 9 bp upstream of a predicted MFS efflux pump, within 2 bases of a previously reported mutation selected during evolution of CIP resistance in *A. baumannii* in planktonic conditions^34^. Consistent with MFS transporter over-production, we found that the *mfsA*(-9) mutation increased the *mfsA* mRNA levels by ∼9-fold relative to the parental strain, based on q-rtPCR analysis (Fig. 4B). Significantly, when the *mfsA*(-9) mutation was combined with the *adeL*P131H mutation, the CIP MIC increased to ∼4ug/ml approaching that of the quadruple mutant strain isolated from mouse passage 15 (Fig. 4C; Table S1). A single nucleotide change located nearby the *mfsA*(-9) mutation that arose transiently in the immunodepleted Lineage 3 (Fig. 2D; *mfsA*(-19)) behaved nearly identically (Figs. 4B,C). Therefore, combining two mutations that resulted in upregulation of both the AdeFGH pump and a putative MFS transporter was sufficient to provide CIP resistance above the clinical breakpoint, approaching that seen for the quadruple mutant.

### A hotspot for mutations within *lpxD* is tightly linked to fluoroquinolone resistance in clinical isolates

The mutation *lpxD* T118_A119insA (locus tag: ACX60_07955; GenBank: AKQ26661.1) was consistently found in CIP^R^ isolates from mice. This alteration within an acyltransferase involved in lipooligosaccharide (LOS) biosynthesis^35–37^ showed no effect on MIC (Table S1). To test for clinical linkage of *lpxD* mutations to drug resistance, we downloaded the genomes of all 8666 *A. baumannii* genomes clinical isolates from the PATRIC database^38^. Focusing on nonrepetitive sequences that had antibiotic resistance profiles, 1830 were resistant to ciprofloxacin and 215 were susceptible. Of the CIP^R^ isolates, 883 genomes had an E->K change at residue number 117, with a *z*-score ≥ 18 (Supp. Fig. 1). In comparison, only 9 of 215 CIP^S^ isolates had the same mutation. By Fisher’s exact t test, p<0.0001, supporting the siginificance of these observations.

To determine if the overrepresentation of the E117K allele could be due to outgrowth of a single clone, we performed multi-locus sequence typing (MLST) using the Pasteur *A. baumannii* scheme, generating 7-gene allele profiles ^39^. Of the 123 distinct MLST groups, 16 had both CIP^R^ and CIP^S^ isolates (Supp. Dataset 1; Supp. Fig. 2), although over half of the CIP^R^ isolates belonged to two MLST types (ST2 and ST3). To avoid skewing the results due to this overrepresentation, we limited the number of isolates per ST group to 1, reducing strains analyzed to 139 across 123 distinct MLST groups (Fig 5A), including 59 CIP^R^ isolates and 80 CIP^S^. Even after limiting the clonal effects in this fashion, we still observed overrepresentation, with the E117K allele found in 13 of 59 CIP^R^ isolates (Fig. 5B), while only 3 of 80 CIP^S^ isolates had the same alterations (Fig. 5B; p= 0.0011, Fisher’s exact t test). Both a whole genome phylogenetic tree (Supp. Fig. 3) and clustering analysis of the 7-gene allelic profiles from these 139 clinical isolates (Fig. 5C) identified CIP^R^ isolates that shared parents with CIP^S^ ones, further arguing against clonal effects for the enrichment.

**Figure 5.**
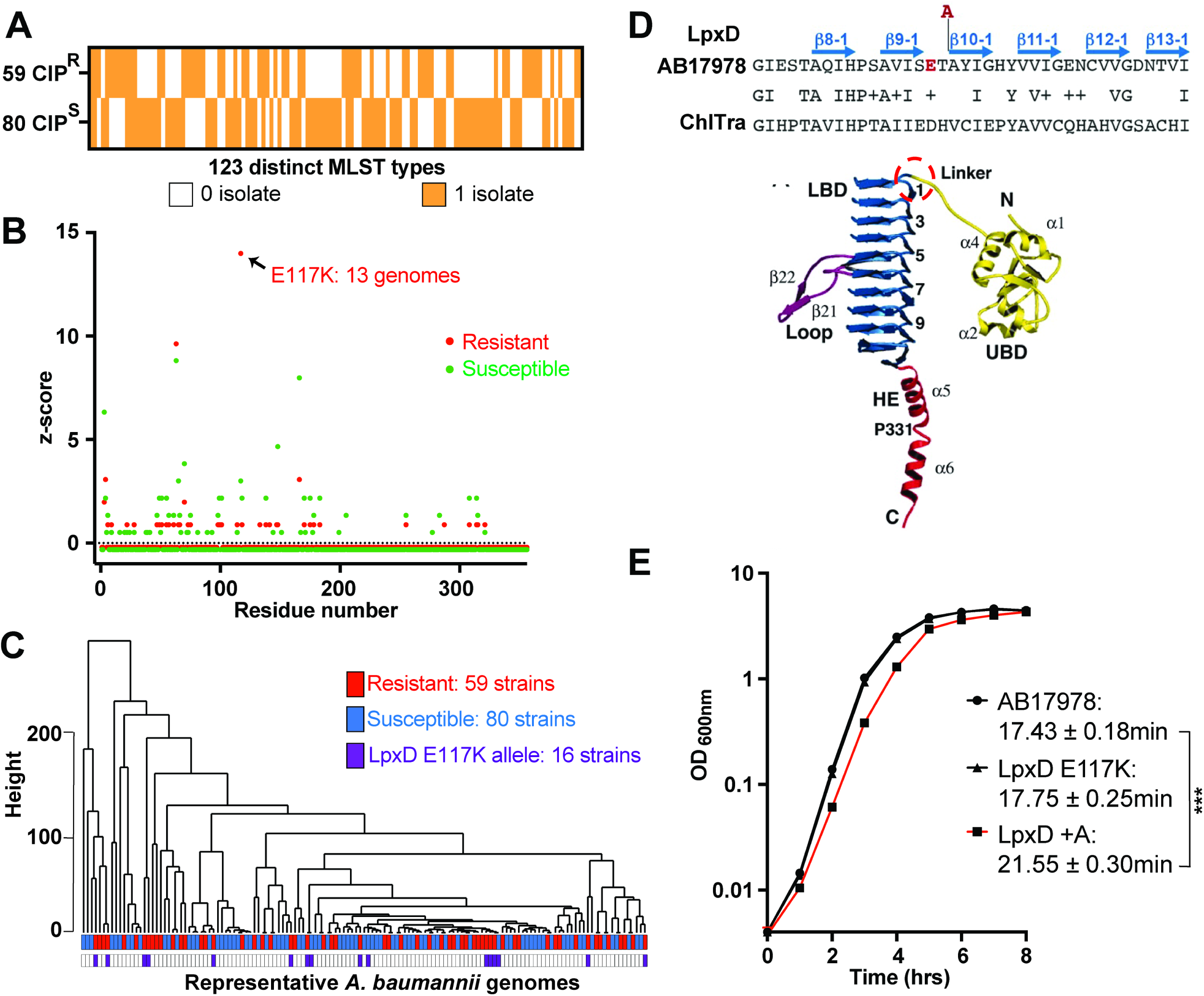
The LpxD E117K allele is tightly linked to clinical fluoroquinolone resistance. A) Assembled genomes of 139 clinical isolates with ciprofloxacin resistance profiles from PATRIC database^80^. Displayed is heatmap showing the presence (orange; 1 isolate) or absence (white; 0 isolate) of clinical isolates for each ST type per each resistance group. B). LpxD sequences from resistant (red dots) and susceptible isolates (green dots) were aligned and compared to ATCC 17978 and the total number of variants per residue was calculated for each group of genomes. The total number of amino acid changes was normalized and presented as a *z*-score for each group, plotted against the LpxD residue number. C). Hierarchical clustering of the 7-gene MLST profile from 139 genes showing Euclidean distance as height and nodes color-coded with resistance profile (red: CIP^R^; blue: CIP^S^) and the presence of LpxD E117K allele in purple. D). Structure of *Chlamydia trachomatis* LpxD protein^38^ showing the presumed site of the *A. baumannii* E117 based on sequence similarity. E). Broth growth of AB17978 strains differing by single changes in *lpxD*. Shown are strains harboring either the WT, *lpxD* T118A119_insA or *lpxD* E117K alleles. Statistical significance was performed using one-way ANOVA followed by Dunnett’s multiple comparison. ***: p-value < 0.001 for LpxD + A strain against WT. Data shown are mean ± SEM.

Interestingly, the alteration at residue E117 (z-score: 8.98; p-value, < 0.00001) directly proceeds the T118 allele identified in our mouse experiments, with both variants predicted to be in a turn between two beta sheets (Fig. 5D)^38^. Although the significance of this linkage is unclear, the fact that the *lpxD* T118_A119insA has a fitness defect in broth (Fig. 5E), raises the possibility that mutations in this turn could contribute to a persistence phenomenon in tissues leading to evolution of drug resistance.

### Evolutionary replay experiments reconstruct the pathway to drug resistance

The observation that resistant organisms outcompeted a persistent strain (Fig. 2D) could be due to a coincidence that occurred during serial passage or due to antibiotic-driven selection for strains with increased MIC over persistent strains. We performed three evolutionary replay experiments^40^ with neutrophil-depleted Lineage 3 variants to distinguish between these possibilities. In the first (Fig. 6A), bacteria were harvested after passage 9 and cycled 3 times in duplicate in neutrophil-depleted animals to test if the AdeL ΔI335A336 allele, a 6 nucleotide in-frame deletion identified in cyclophosphamide treated animals, would again be outcompeted by low abundance drug-resistant mutants (Fig. 2D). The second approach was to passage three times a mix of a colony-purified drug persistent AdeL ΔI335A336 single mutant (P9Cy3.1; Table S1) with a colony-purified drug resistant quadruple mutant (Cy31; Table S1) in the approximate ratio present at passage 9 (95:5; Fig. 6B). The third was to test the model that neutrophils restrict the outgrowth of strains having activating mutations in *adeL* (Fig. 6C).

**Figure 6.**
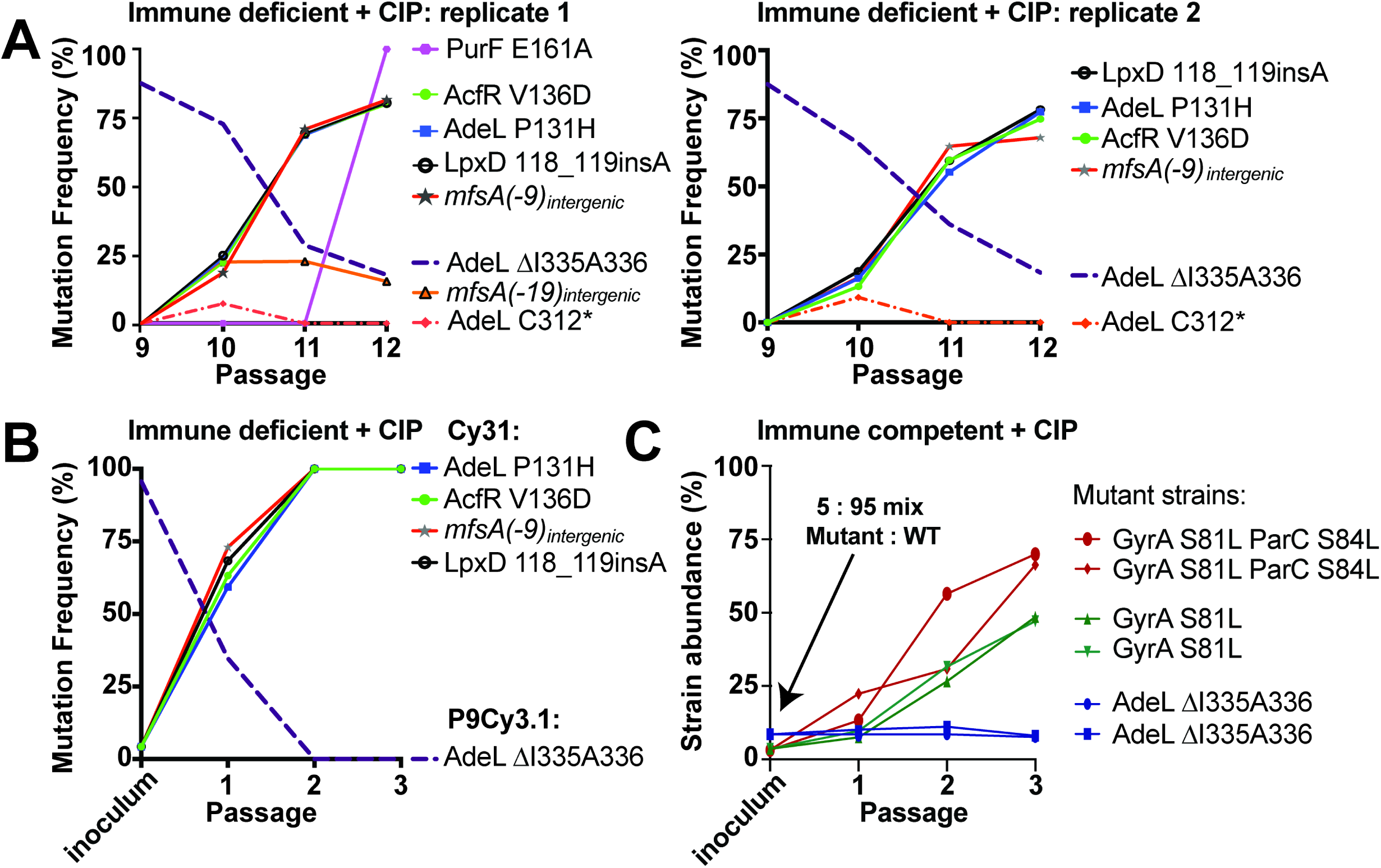
The evolutionary trajectories of isolates acquired throughout passaging in neutrophil-depleted mice is due to fitness advantage in presence of drug. (A) A single *adeL* mutant is outcompeted during infection of CIP-treated mice by a quadruple mutant (Cy31) having drug resistant *adeLmfsA* alleles. The passage 9 pool was oropharyngeally inoculated into neutrophil-depleted mice in duplicate but separate lineages and passaged three times in the presence of CIP treatment, subjecting each passage to deep sequencing. A-B)Displayed are plots showing relative abundance of all mutations identified in deep sequencing pools of the duplicate passaging experiments as a function of the passage number) Competition of purified *adeL* and quadruple mutant having *adeLmfsA* alleles in presence of CIP results in predominance of the quadruple mutant. The two strains were mixed at a ratio 95:5 (single : quadruple mutant), and oropharyngeally inoculated into neutrophil-depleted mice in the presence of CIP treatment through three successive passages. Deep sequencing of the initial inoculum and each subsequent passage was performed to determine the ratio of the two strains, and the relative abundance of each mutation is displayed as a function of passage. C) The presence of neutrophils prevents enrichment of an *adeL* mutant during CIP challenge in the mouse. Competition of designated mutants (*adeL*ΔI335I336, *gyrA*S81L, or *gyrA*S81L *parC*S84L) against wild type AB17978 was performed. The strains were mixed at a ratio 95 : 5 (AB17978 : mutant), oropharyngeally inoculated into immunocompetent mice. and then passaged in duplicate but separate lineages three times in the presence of CIP treatment. The abundance of each mutant was determined by measuring the colony forming efficiency (CFE) at an antibiotic concentration that was below mutant MIC but above that of wild type AB17978 (AdeL ΔI335I336 MIC: 0.75 µg/mL, GyrA S81L MIC: 0.75 µg/mL, or GyrA S81L + ParC S84L MIC: 2 µg/mL), and the CFE versus passage number was displayed and expressed as strain abundance (%). CFE = number of colonies on antibiotic-containing medium / number of colonies in the absence of antibiotic.

When the *adeL* mutant was analyzed in competition within the pool (Fig. 6A) or when mixed 95:5 with colony purified quadruple mutant (Fig. 6B), we observed an increase in bacterial burden and fraction of bacteria showing CIP MIC >2 after passaging the pools in neutrophil-depleted CIP-treated mice (Supp. Fig. 4). Whole genome sequencing of the bacterial populations harvested after each passage revealed a near identical evolutionary trajectory as seen from the original experiment, with the single mutant associated with drug persistence eventually outcompeted by drug resistant mutants (Fig. 6). Furthermore, in one of our replicate Lineages (Fig. 6A), another mutation located in the intergenic region between *rpmL* and *mfsA* (*mfsA*(-19)) arose that had appeared transiently in immunodepleted Lineage 3 (Fig. 2D), further arguing for the contribution of this regulatory region in resistance evolution. Therefore, these two reconstructions (Figs. 6A,B) demonstrate that the evolutionary trajectory is reproducible, and can be regenerated by phenotypically similar mutations that arise spontaneously.

In the third experiment, single *adeL* mutants were unable to overgrow the WT in the presence of CIP in immunocompetent animals, in contrast to what was observed in passage 9 of the neutrophil-depleted animals. When a 5:95 (*adeL*:WT) mixture was passaged in immunocompetent mice, the mutant was unable to increase its population share (Fig. 6C), mimicking the failure of *adeL* activating mutations to overgrow the pool in any of the immunocompetent Lineages (Fig. 2A). In contrast, CIP^R^ mutants having canonical resistance alleles in *gyrA* or *gyrAparC* were able to compete efficiently with WT in the presence of neutrophils, with the GyrA(S81L) ParC(S84L) double mutant growing from 5% to 70% of the population within 3 passages (Fig. 6C). This argues that depleting the immune response allows the outgrowth of persistence mutants, facilitating complex pathways to drug resistance that are blocked in immunocompetent hosts.

### Ciprofloxacin treatment within the host simultaneously selects for increased resistance and enhanced fitness

Antibiotic resistance is often associated with a fitness cost^41^ that may be compensated over time by continued selection of secondary genotypes that overcome these costs. The inability of the *adeL* mutant to overgrow WT during infection in the presence of CIP in immunocompetent animals may reflect this point. Mutants with high resistance, however, could maintain their selective advantage over non or low-resistance mutants even in the absence of drug treatment. To test this model, the relative fitness of the drug resistant mutants derived from neutrophil-depleted Lineages was compared to the parent strain AB17978. In the absence of CIP, variants isolated after passage in neutrophil-depleted mice showed varying degrees of subtle growth impairment in broth relative to the parental strain (Fig. 7A; Cy11, p< 0.01). During animal infections in the absence of antibiotic, these growth defects were amplified. When immunocompetent mice were challenged with the high fitness *gyrAparC* mutant in competition with the WT, the two strains showed similar levels of colonization (Fig. 7B). In contrast, each of the drug resistant mutants derived from neutrophil-depleted mice competed poorly in immunocompetent animals (Fig. 7B; p <0.0001).

**Figure 7.**
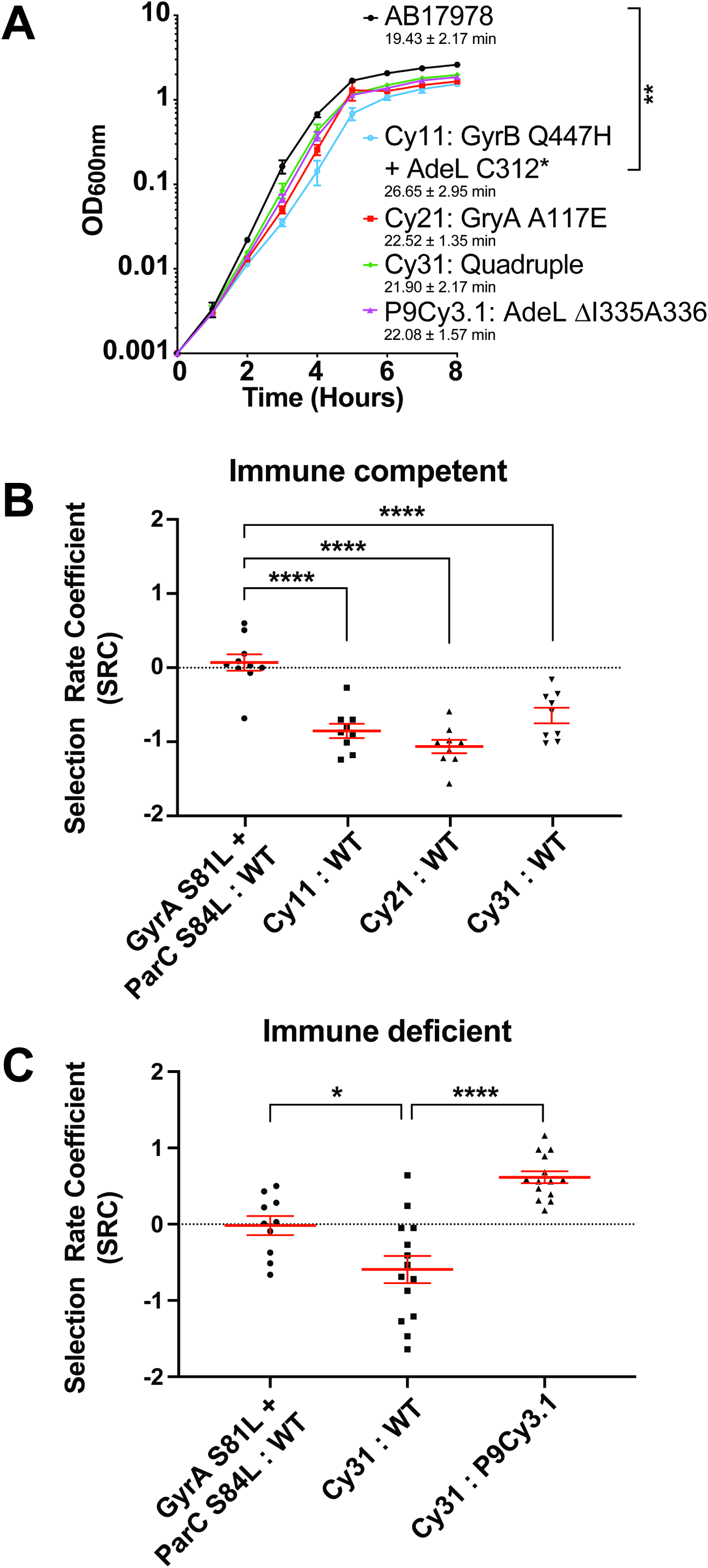
Simultaneous selection for increased resistance and fitness. A) Growth rates in broth culture of noted isolates from passages in mice. Significance determined by one-way ANOVA followed by Dunnett’s multiple comparison: **: p<0.01 for Cy11 against WT. C312*: mutation of Cysteine to stop codon at residue 312. B) Neutrophil-replete and C) Neutrophil depleted (cyclophosphamide treated) BALB/C mice were challenged oropharyngeally with noted strain combinations for 24-hours to determine the selection rate for each competing pair of strains. Selection rate coefficient (SRC) was determine by SRC = LN(Strain 1 output/input ratio) – LN(Strain 2 output/input ratio)(Materials and Methods) ^72^. Data are mean ± SEM, with significance determine by one-way ANOVA followed by B) Dunnett’s or C) Tukey’s Multiple Comparison. *p<0.05; ****p<0.0001.

Although the lowered fitness of mutants in immunocompetent animals further supports the model that outgrowth of unique resistance alleles occurs in the absence of neutrophils, it did not directly test the model that selection in neutropenic mice increased fitness relative to first-step CIP^R^ variants. In the absence of CIP treatment, the parent strain showed a subtle advantage over the Lineage 3 quadruple mutant isolate Cy31 (Table S1) under neutrophil-depleted conditions, although the results were clearly not as robust as in the presence of neutrophils due to increased scatter (Fig. 7C; Cy31 against WT). More striking is the fact that the Cy31 strain had a clear fitness advantage over its predecessor, the Lineage 3 *adeL* mutant, when competed in the absence of antibiotics (Fig. 7C; Cy31 against P9Cy3.1). Therefore, in competition with persister mutants, ciprofloxacin simultaneously selected for an isolate with increased CIP resistance and enhanced fitness relative to mutant competitors. This predicts that drug-resistant mutants were able to evolve to heightened fitness to replace persister mutants. As fitness increases, resistant mutants should be able to compete with sensitive strains even after cessation of antibiotic treatment, potentially escalating treatment failure and providing fertile ground for the emergence of resistant strains with high fitness.

## Discussion

Comparative analyses of bacterial pathogen databases indicate that clinical antibiotic resistance is associated with mutations located outside well-characterized drug targets ^20–22^. Analysis of clinical resistance is often retrospective, with a few exceptional studies that have allowed identification of a timeline of bacterial resistance in human populations^28–32, 42–44^. Our study bridges an important gap in understanding how resistance evolves, by following experimental evolution of fluoroquinolone resistance in a pneumonia disease model. Strikingly, in the absence of one arm of immune pressure, *A. baumannii* resistance arose after few passages in murine lungs during CIP treatment (Fig. 1B), resulting in treatment failure. Conversely, the overall frequency remained low in immunocompetent mice, and this did not lead to treatment failure (Fig. 1B).

The absence of neutrophils resulted in the identification of rare alleles associated with drug resistance. In contrast to the canonical target site GyrAS81LparCS84L mutation which shows high fitness under conditions tested, the *gyrA*A117E and *gyrB*Q447H alleles rarely observed in clinical strains predominated in two of the lineages from neutrophil-depleted animals (Fig 2E). These unusual isolates showed fitness defects during disease and/or in culture (Fig. 7) that make them unlikely to arise in the presence of intact immune functions, consistent with evolutionary trajectories being determined by the immunological state of the host. Therefore, we propose that immunocompromised hosts are incubators for the generation of unique drug resistant variants, with uncertain outcomes after pathogen exposure to the community at large. Although the variants we described have reduced fitness relative to WT, continued passage demonstrated that stepwise increase in drug resistance was associated with stepwise increase in fitness (Fig. 7). Similar fitness effects have been obtained during continued passage in culture^34^. The recent demonstration of enhanced viral evolution of SARS-CoV-2 in an immunocompromised patient undergoing therapeutic antibody treatment is a graphic example of the potential interplay between immunity, antimicrobials and evolution in the clinic, with potential largescale community effects^45^. Furthermore, persistence mutants have been demonstrated to arise during continued antibiotic therapy in an immunocompromised patient treated for vancomycin-resistant enterococcus^46^, tying the results presented here with documented clinical outcomes.

Antibiotics primarily act by targeting essential cellular functions or pathways ^47, 48^. In response to transient antibiotic exposure, a proportion of bacteria typically persist or tolerate this treatment with mutations arising that can increase the fraction of the surviving population^49–51^. The mechanisms for allowing persistence/tolerance likely differ between bacterial species and vary among antibiotics^52^, but it is generally agreed that persistence/tolerance is caused by cell dormancy^53–55^ or transient expression of efflux pumps and stress response pathways^28, 56^. Persistence/tolerance mechanisms may play an important role in the relapse of bacterial infections ^57–59^. In our study, we found that persistence in *A. baumannii* can be promoted by upregulation of a drug efflux pump, AdeFGH ^26^. It is known that clinical isolates often overproduce efflux pumps that remove multiple antibiotics from the bacterial cytoplasm^26, 60^, and that antibiotic persistence is associated with such pump upregulation^61, 62^. Of note, mutations upregulating this pump are rarely isolated in the laboratory through simple antibiotic single-step selections on solid agar, indicating there may be a special environment in the lung that allows outgrowth of these mutants.

There are three important repercussions from the analysis of the AdeL efflux overproducer mutations. First, overproduction results in drug treatment failure during pneumonic disease in the mouse, in spite of the fact that these bacterial strains are CIP-sensitive based on the international standard for clinical breakpoints^27^. Therefore, clinical drug treatment failure in the absence of identified antibiotic resistance^9^ could be explained by the outgrowth of drug persistent mutants, particularly in the immunocompromised host. Second, the *adeL* mutations provide a molecular basis for *A. baumannii* to develop second-step variants that lead to CIP resistance above the clinical breakpoint. Of interest, inhibitors of efflux pumps have been suggested to re-sensitize bacteria to multiple antibiotics^63^, but perhaps such a regimen may also delay progression to resistance. Finally, isolates from drug treatment failure occurring in immunocompromised patients could generate a pool of precursor mutants giving rise to resistant isolates that eventually infect a broad range of patients. A number of studies have similarly argued that efflux pump mutations provide a critical pool for the multi-step evolution of antibiotic resistance ^33, 64, 65^

We propose a model of bacterial resistance progression in immunodepleted hosts (Fig. 8). When infecting a host, the antibiotic susceptible bacteria colonize at a relatively low rate. With active transmission and continuous CIP treatment, there is outgrowth of bacterial persisters that have MICs below the clinical breakpoint, increasing the bacterial load at the infection site. On continued treatment, or transfer to a similar host undergoing CIP treatment, fully resistant mutants can then outgrow the population. The presence of intact immune function interferes with outgrowth of these mutants, but the identification of persister mutants in the presence of immune function indicates that such alleles could act as enablers of drug resistance in all hosts, particularly under conditions of temporary breaching or depression of innate immunity.

**Figure 8.**
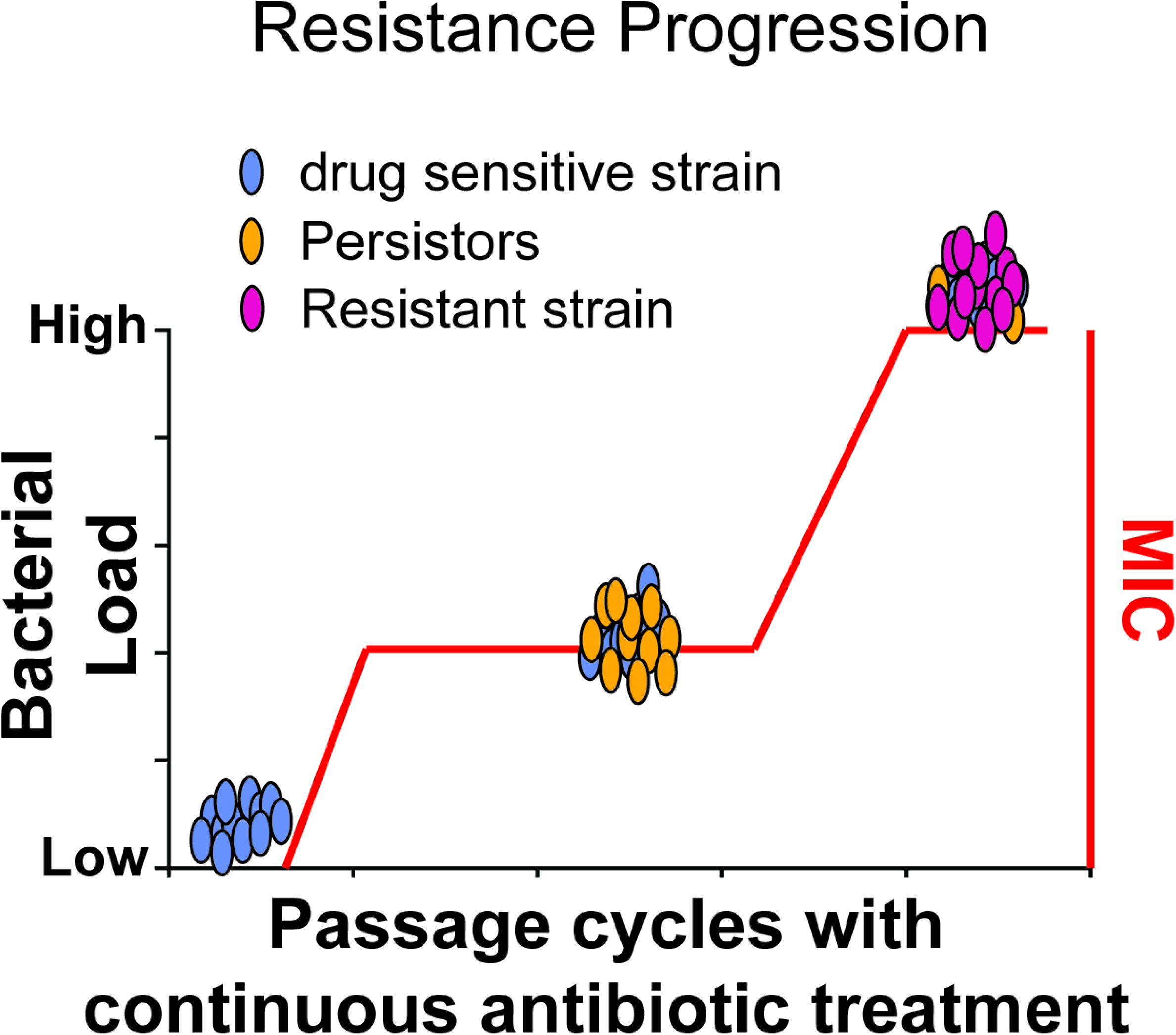
Model of bacterial resistance progression. The antibiotic susceptible inoculum (blue) colonizes poorly in presence of antibiotic. During continued passage with CIP, in the absence of neutrophils, the persister mutants (yellow) arise as first step mutations showing intermediate MICs. With continued passage under the same antibiotic treatment regime, second step mutations arise that have increased fitness in the presence of antibiotic (red) and dominate the population.

Genomic analyses can identify allelic variants linked to drug resistance, but verifying the functional importance of these alleles is hindered by founder effects that are often difficult to discount. The *lpxD* E117K mutation is one such allele associated with resistant *gyrA* S81L- containing genomes. The fact that the adjacent *lpxD* T118 insA allele was isolated during multiple passages in immune-depleted mice argues for a role of these altered residues in supporting the outgrowth of drug resistant organisms. The *lpxD* T118 insA mutation likely alters envelope function. This raises the possibility that the mutation causes subtle changes in permeability that slow drug access to target, or else the slowed growth of strains harboring this allele increases drug tolerance during growth in tissues^52^. Our inability to demonstrate increased tolerance to CIP and the fact that the clinical *lpxD* E117K did not show a fitness defect when grown in LB broth (Fig. 5E) argues that the critical phenotypes may be observed exclusively during growth in tissues, making them difficult to evaluate.

In summary, we hypothesize that resistance progression in clinical isolates follows a similar trend witnessed in our experiments, albeit with greater complexity in the human. Drug sensitive bacteria may colonize both healthy and immune system-compromised patients, but resistance evolution occurs more rapidly within hosts having impaired neutrophil function. The canonical resistance mutations that arise within healthy individuals may compete efficiently with host-evolved mutations in patients having impaired neutrophil function. Hence, the transmission from individuals having intact immune function may pose risks to immunocompromised patients. Adding to the risks subjected to vulnerable patients, we found that *A. baumannii* persistence mutants with MIC levels below the clinical breakpoint were associated with treatment failure, emphasizing the difficulty in treating these patients (Fig. 1B) ^9, 66–69^. While heterogeneity and variability between infected patients is inevitable, this work provides insight into some genetic factors that could predispose individuals to treatment failures in specific clinical contexts. Future work should be focused on how this evolutionary trend towards drug resistance can be controlled, such as tackling the molecular basis for selection of persister or other mutations that lead to treatment failure.

## Material and Methods

### Bacterial Strains

Bacterial strains used in this study are listed in Table S1. *Acinetobacter baumannii* strains are derivatives of ATCC 17978. Bacteria were grown in Lysogeny Broth (LB) (10 g/L tryptone, 5 g/L yeast extract, 10 g/L NaCl) or on LB agar plates (LB supplemented with 15g/L agarose). Broth cultures were grown at 37°C in flasks with orbital shaking at 200 rpm or in tubes with rotation on a roller drum at 56 rpm. Growth was monitored by measuring absorbance spectrophotometrically at 600 nm (A_600nm_). Plates were incubated at 37°C. Antibiotics were used at the following concentrations: Gentamicin: 10 μg/mL (*A. baumannii*) and 50 μg/mL (*E. coli*); Carbenicillin: 100 μg/mL.

### Murine experimental evolution

The *in vivo* passaging experiments were performed identically in either immunocompetent or neutrophil-depleted 6-8 weeks-old female BALB/C mice (Fig. 1A). Neutrophil depletion was induced via cyclophosphamide pretreatment ^70^. Briefly, 150 mg/kg and 100 mg/kg of cyclophosphamide monohydrate (Sigma-Aldrich: C7397) were administered 4 days and 1 day prior to infection, respectively. At the time of infection, mice were transiently anesthetized by inhalation of isoflurane and lung infections were established using ∼10^8^ colony forming units (CFU) of *A. baumannii* from mid-log phase cultures (A_600nm_ ∼0.5) via oropharyngeal aspiration ^71^. For Passage 1, all Lineages were derived from a single broth-grown culture. At 7 and 19 hours post infection (hpi), 100 mg/kg of ciprofloxacin (Sigma-Aldrich: PHR1044) was administered via subcutaneous injection. At 27 hpi, mice were euthanized, and lungs were removed aseptically and homogenized in 1 mL of ice-cold 1x phosphate buffered saline (PBS). Afterwards, the homogenate was plated on large (150 x 15 mm) LB agar plates and incubated for ∼17 hours at 37°C. The total number of bacteria was quantified using CFU. The enriched bacteria were scraped off the LB plates and resuspended in 1x PBS. The resuspension was used for DNA isolation as well as the inoculum for the next passage/round of infection (stored at -80°C in 20% glycerol prior to use). For each round of mouse infection/passage, bacterial frozen stocks were revived by diluting them in LB broth and growing them to exponential phase. Afterwards, ∼10^8^ CFU of the pool was used to establish infection in each mouse. Three separate Lineages of murine infections were maintained in parallel in each condition (immunocompetent or neutrophil depleted), and within each Lineage each starting inoculum was passaged 15 times. Validation experiments involving the reconstruction of identified mutants and re-passaging of these strains in mice were similarly performed.

### Murine Competition Assays

Competition experiments were performed as described previously^71^. Briefly, designated strains were grown to exponential phase (A_600nm_ ∼ 0.5-1.0) and mixed at a 1:1 ratio for the inoculum. Afterwards, ∼10^8^ CFU of the bacterial mixture was used to challenge WT or cyclophosphamide-treated BALB/C mice via the oropharyngeal route as described for the mouse passage experiments. 24 hr post inoculation, lungs were removed aseptically, homogenized and extracts were spread on LB agar plates. To quantify the input and output ratio of each designated strain, both the inoculum and the lung homogenate were serially diluted in 1x PBS and plated on LB agar as well as LB agar supplemented with 2 μg/mL of ciprofloxacin to differentiate between the WT and CIP resistant test strains. Selection rate coefficients (SRC) were determine by SRC = ln(Strain 1 output/input ratio) – ln(Strain 2 output/input ratio)^72^. Prism was used for statistical analysis and significance was determine by one-way ANOVA followed by Dunnett’s or Tukey’s Multiple Comparison. *p<0.05; ****p<0.0001.

### Evolutionary replay experiments

For the evolutionary replay experiments (6A), the saved Cy3 Passage 9 pool was used as inoculum. For the evolution experiments involving isolates (Figs. 6B and 6C), designated strains were grown to exponential phase and mixed at a 5:95 ratio in the inoculum. In both cases the inoculum was administered via the oropharyngeal route to mice and passaged three times in the presence of CIP treatment in a near identical manner as the initial passaging experiments. Each pool/mixture was performed in duplicate. The bacterial pools after each passage were saved, and variant frequencies at each passage were determined as described above. The abundance of each mutant in the isolate mixtures was determined by measuring the colony forming efficiency (CFE) at an antibiotic concentration that was below the mutant MIC but above that of wild type AB17978. The CFE versus passage number was displayed and expressed as strain abundance (%). CFE = number of colonies on antibiotic-containing medium / number of colonies in the absence of antibiotic.

### Determining the fraction of resistance as a function of ciprofloxacin concentrations in each pool

To evaluate the fraction of bacteria acquired from each passage that are viable on culturing in various concentration of ciprofloxacin, the glycerol stock of each pool was revived in fresh LB broth and grown to exponential phase. A total of ∼10^7^ CFU were used for serial dilutions in 1x PBS and 10 μl from each diluted culture was spotted on LB agar plates containing the following concentrations of ciprofloxacin: 0, 0.25, 0.5, 1, 2, 4 and 8 μg/ml. After an overnight incubation at 37°C, colonies were counted, and CFU/ml was calculated. A detection limit of 100 CFU/ml was used. The fraction bacteria resistant to a certain concentration (C_0_; C_0_ > 0) of ciprofloxacin is calculated as (CFU/ml at C_0_ - CFU/ml at all concentrations above C_0_) / Total CFU. A stacked bar plot of the data was generated using Prism GraphPad (Fig. 2).

### Whole Genome Sequencing

Genomic DNA (gDNA) was extracted from bacteria using the DNeasy Blood and Tissue kit (Qiagen). Library preparation was performed using Illumina Nextera with previously described modifications ^73^. Libraries were enriched with 8 cycles of PCR and sequenced using HiSeq2500 at Tufts University Core Facility using single-end 100 bp reads. Reads were aligned to the *A. baumannii* ATCC 17978 genome and its plasmids (GenBank Accession: CP012004; pAB1: CP000522; pAB2: CP00523; pAB3: CP012005), and variants were identified at 5% cutoff using breseq 0.32 ^74^. Reads were aligned to *A. baumannii* ATCC 17978 reference genome (https://genomes.atcc.org/genomes/e1d18ea4273549a0) to include an additional 44kb gene cluster ^75^. The identified mutations are listed in Supplemental Dataset 2. The predicted functional impact of substitution variants was determined by using PROVEAN ^76^.

### Persistence Assay

*A. baumannii* was grown in 3 individual LB broth cultures overnight from separate colonies. The cultures were diluted 1000-fold into 8 ml LB broth and incubated by rotation at 37°C for 2-3 hours until reaching mid or late exponential growth (A_600nm_ ∼ 0.3-0.8). Ciprofloxacin was added to reach a final concentration of 10 μg/ml, the concentration regarded as 20x the MIC for the parent ATCC 17978 *A. baumannii* strain. These cultures were then incubated at 37°C for up to 24 hours. After 0, 1, 3, 6, and 24 hours, 500 μl of culture was removed, washed twice in 1x PBS, resuspended in 500 μl PBS, and sequentially diluted in 1x PBS. Serial dilutions were used for CFU quantification by spot plating on LB agar plates. Dilution factors harboring quantities ranging from 3-35 CFUs were used to calculate the CFU/ml. The limit of detection by this assay was determined to be 375 CFUs. The CFU/mL across the biological replicates at each time point throughout the drug challenge were plotted as mean ± Standard Error of Mean.

### qRT-PCR gene expression analysis

Bacteria were grown to early stationary phase in LB broth, RNA was harvested and purified using RNeasy kit (Qiagen) followed by cDNA synthesis using SuperScript VILO cDNA kit (Thermo Fisher). The cDNA was then amplified with PowerUp SYBR Green Master Mix (Applied Biosystems) via a StepOnePlus Real-Time PCR system (Applied Biosystems) per manufacturer’s instructions, and target amplification efficiency was evaluated by generating a standard curve with dilutions of cDNA (>95% amplification efficiency for each primer pair). Primers were designed to amplify regions of around 150 bp internal to genes (Table S2). Triple technical replicates were examined per biological sample and at least three biological replicates per strain were tested, with controls lacking reverse-transcriptase included to verify a lack of contaminant genomic DNA. Transcript levels of specific targets from each strain were evaluated by the comparative 2^-ΔΔCt^ method to the parental strain, normalizing to that of the endogenous control 16s. The transcript level for each target across biological replicates was plotted as mean ± Standard Error of Mean (Fig. 3&4).

### MIC determination

Minimum inhibitory concentrations (MICs) were determined by broth microdilution. Overnight cultures of strains of interest were diluted 1000X in fresh LB broth and grown to mid-logarithmic phase (A_600_ ∼0.5). Afterwards, cultures were diluted to a final A_600_ = 0.003 and tested in the presence of two-fold dilutions of antibiotics. 200 μl of culture-antibiotic mixture was then aliquoted to a 96 well plate (COSTAR) and technical duplicates were performed in a Biotek plate reader with rotation. Growth was monitored by measuring A_600_ at 15 min. intervals for 16 hr, and the MIC was determined as the lowest concentration of drug that prevented growth, using at least three biological replicates for each strain. The MICs across biological replicates were plotted as mean ± Standard Error of Mean (Fig. 4).

### Molecular cloning and mutant construction

Plasmids and primers used in this study are listed in Table S2. The mutant strains were constructed through sequential cloning into pUC18^77^ then pJB4648^78^ as previously described^13^. Briefly, the mutant allele from each strain of interest was amplified alongside upstream and downstream segments to generate a PCR product ∼1500 bp in length flanked by appropriate restriction sites. The PCR product was then ligated into pUC18 and propagated in *E. coli* DH5α. The PCR product was then subcloned from pUC18 into pJB4648 and propagated in *E. coli* DH5α λpir. After sequence confirmation, the pJB4648 plasmid construct containing the desired mutation was introduced into *A. baumannii* via electroporation. Markerless, in-frame mutations were isolated via homologous recombination as described ^13^.

### Comparative sequence analysis

Genome sequences from 8666 *A. baumannii* clinical isolates and ciprofloxacin resistance profiles for 2618 isolates were downloaded from PATRIC database in Nov 2021 as described in results. MLST analysis was performed using Pasteur scheme^39^ with the publicly available tool “mlst” (https://github.com/tseemann/mlst). The LpxD protein sequence from ATCC 17978 was compared to these 2045 isolates using tblastn to identify the amino acid locations of mismatches and gaps. The nucleotide sequences with the best matches to the LpxD protein were extracted and translated into protein sequences. For each isolate, the translated LpxD sequence was compared to the reference LpxD sequence from AB17978. The nonsynonymous changes were recorded for each clinical isolate. The nonsynonymous changes at each residue were then summarized to reflect the total number of clinical isolates showing those changes. The raw numbers were further transformed into *z*-scores to determine if a particular residue was significantly overrepresented by nonsynonymous changes linked to CIP^R^ or CIP^S^ phenotypes (Supp. Fig. 1). In order to reduce clonal effects, one clinical isolate per ST group was selected randomly. Multiple aligned LpxD protein sequences are shown in Supp. Dataset 3. One isolate had no hit for LpxD. The MLST profiles from 7 housekeeping genes were used to calculate the Euclidean distance, and hierarchical clustering was determined using R (Fig. 5A). A phylogenetic tree from these 139 genomes was built using mashtree ^79^ and visualized in Geneious Prime (https://www.geneious.com/) (Supp. Fig. 3). The presence and absence of isolates was summarized and visualized in a heatmap using Prism GraphPad v9 (Fig. 5B; Supp. Fig. 2). Within each group, the mismatches/gaps at each amino acid location within LpxD were displayed as number of genomes having mismatches/gaps at that location. A table of the data was constructed showing the specific amino acid in the rows, the genome groups harboring these mutations in the columns, and the total number of mismatches/gaps in the cells. The results in each cell were then normalized using *z*-scores with the mean and standard deviation from individual genome groups. The *z*-score was plotted as function of each residue.

### Calculation of doubling time in broth culture

Overnight cultures of bacteria were diluted 1000-fold in fresh LB broth and grown for 8 hours. The A_600_ values were measured every hour using a spectrophotometer, and the log2 transformed values were plotted as a function of time (hr). To calculate the doubling time, a modified sigmoid function 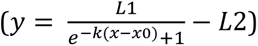 was fitted to the plotted curve (y: log2 transformed A_600_ value, x: time in hrs, x_0_: timepoint when the growth rate was the fastest, k, L1, L2: rate constants), with the fastest growth rate for each culture determined as the time in which second derivative of the fitted curve approached 0 (or when ), at which point the doubling time was represented as the inverse of the slope 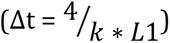 (Code available at: https://github.com/huoww07/calulate_bacteria_doubling_time). To evaluate the goodness-of-fit, the R^2^ value was calculated for each growth curve assay (the R^2^ ranges from 0 to 1, representing the worst to the best fit). At least three biological replicates were performed for each test strain (Table S3, Table S4) and the average doubling time and standard deviation was calculated. Statistical significance was performed using one-way ANOVA followed by Dunnett’s multiple comparisons.

## Accession numbers

The datasets generated during the current study are available in the Sequence Read Archive (PRJNA485355). Detailed accession numbers for each sample are listed in Table 1.

## Acknowledgements

This work is supported by NIAID U01AI12430 to TvO, JWR, VSC and RI, NIH R21AI128328 to RI and T32AI07422 support for JH-B. We thank the colleagues in all of our labs for support and constant critiquing of the experimental strategies used here. We thank all anonymous reviewers for critical comments and suggestions.

## Author Contributions

RI, JR and TvO devised the study. LB, JH and WH performed the mouse experiments. WH, EH and JH performed the wet-lab experiments. WH and LB performed the whole genome sequencing and bioinformatics analyses. CM, VC, EG and TvO collaborated on the bioinformatics analyses and performed training. WH, LM, EH, JH and RI interpreted the results and wrote the manuscript. WH, LB, EH, JH, CM, EG, VC, TvO, JR and RI approved the final manuscript.

## Figure Legends

**Supplemental Figure 1.**
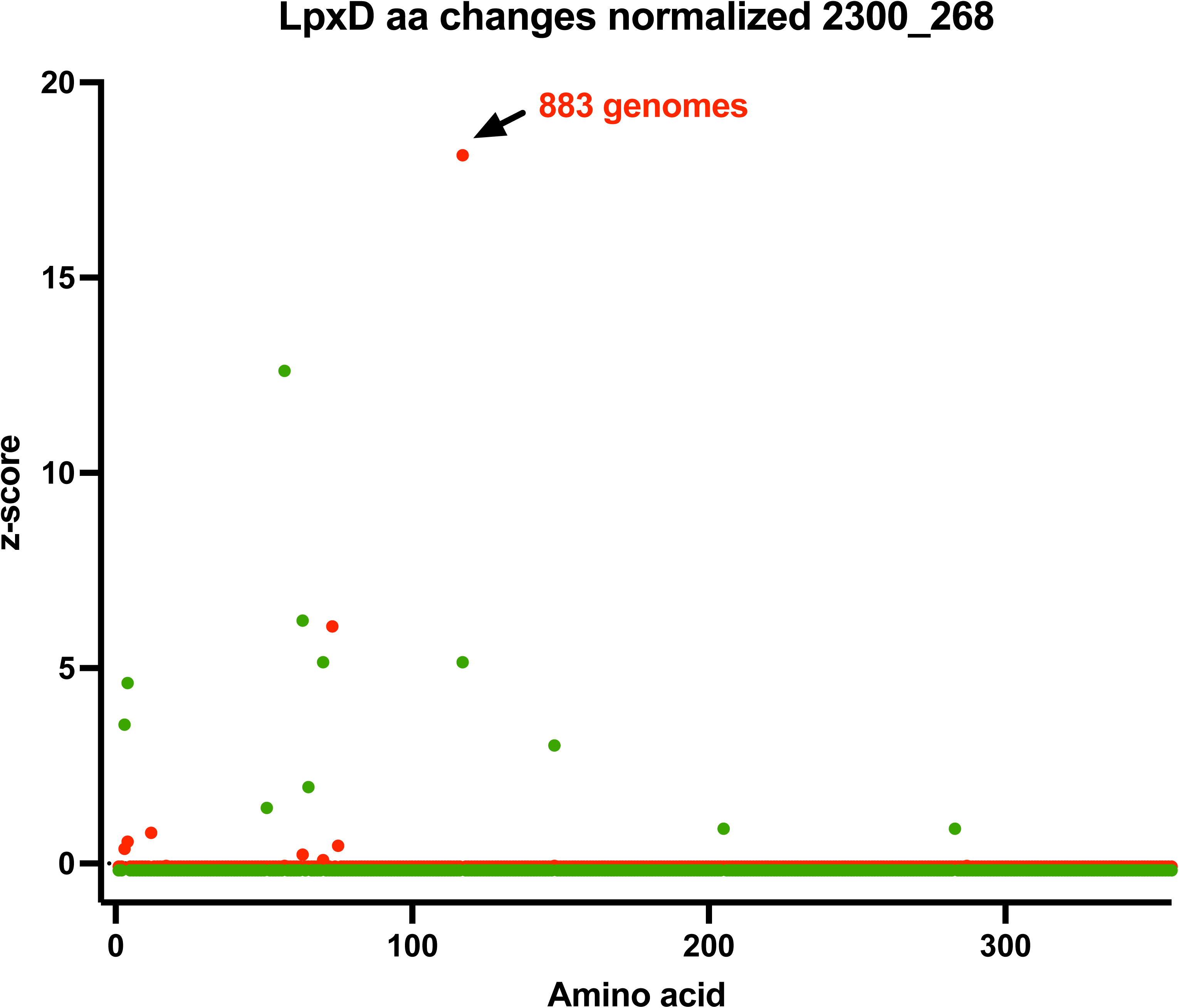
The LpxD E117K allele is tightly linked to clinical fluoroquinolone resistance. Assembled genomes of 2045 clinical isolates were obtained from PATRIC database^80^. LpxD sequences from all genomes were aligned and compared to ATCC 17978 and the total number of variants per defined amino acid was calculated for each group of genomes. The total number of amino acid changes was normalized and presented as a z-score for each of the three groups, plotted against the residue numbers.

**Supplemental Figure 2.**
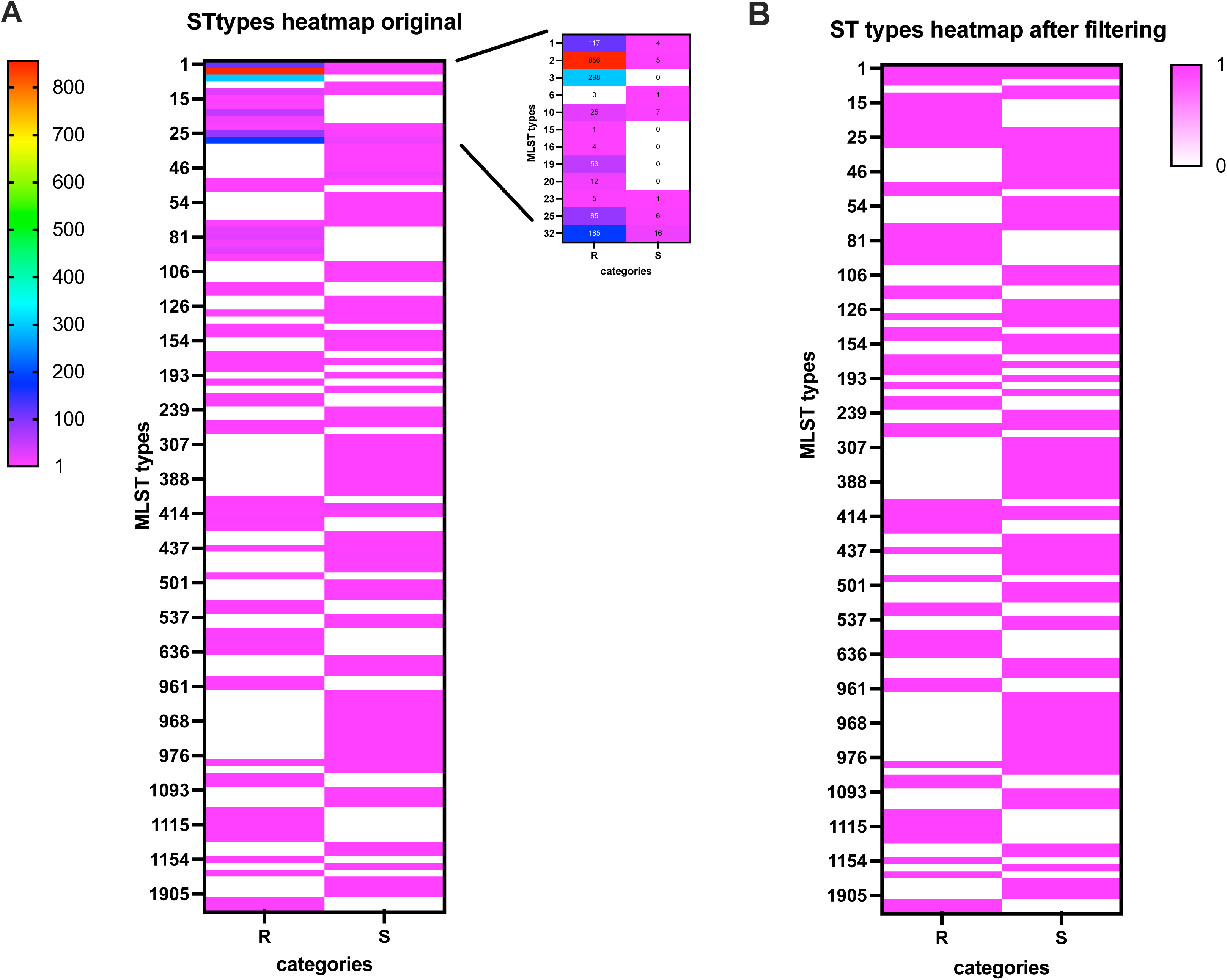
Distribution of MLST type among select clinical isolates. A) Heatmap showing MLST enrichment among 1830 genomes from ciprofloxacin resistant isolates and 269 genomes from susceptible isolates. In order to reduce the populational clonal effect, analysis was limited to only one representative genome per ST group. Heatmap in B) shows the distribution of MLST types after filtering.

**Supplemental Figure 3.**
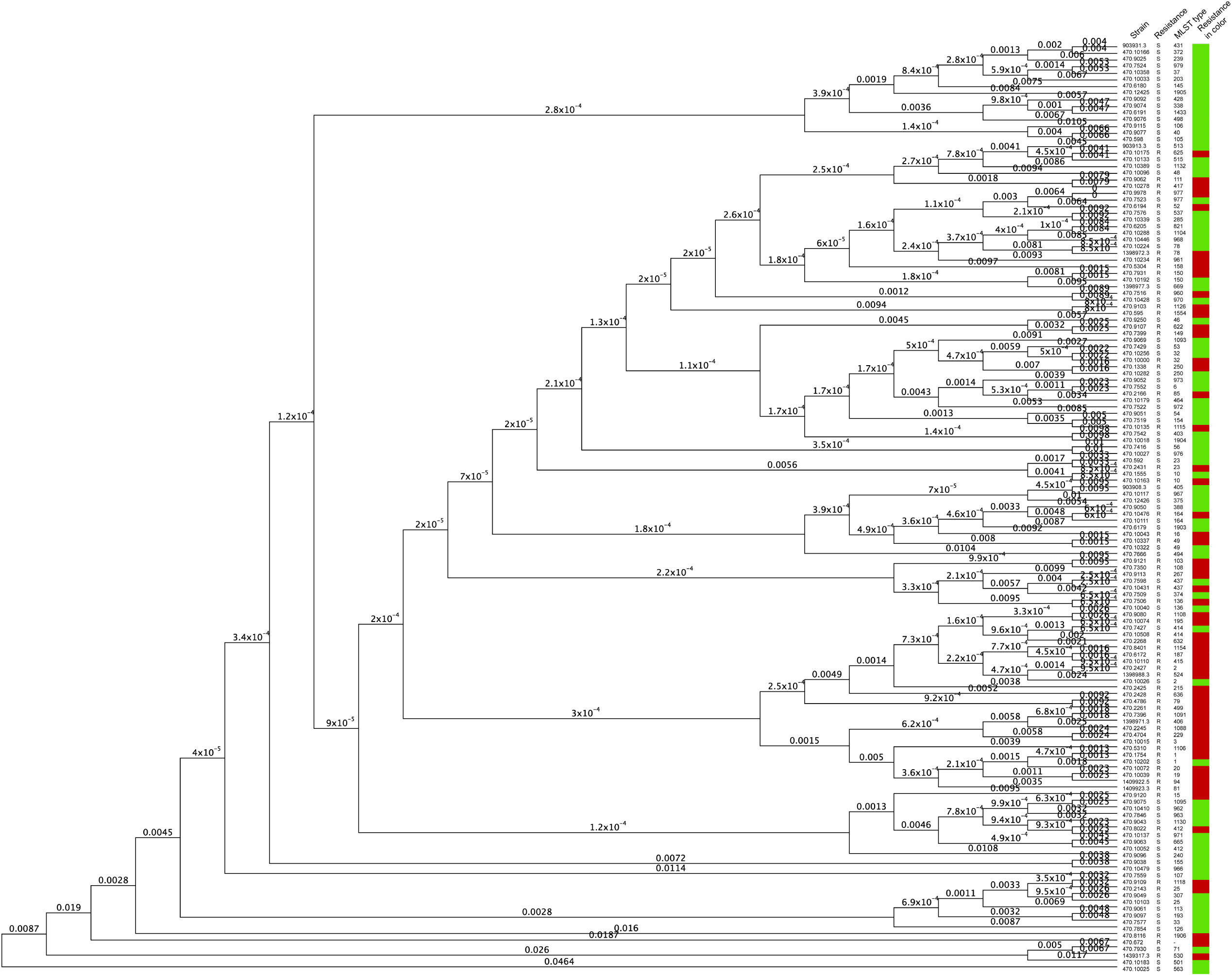
Phylogenetic tree of genomes from 139 clinical isolates. The genome contigs were analyzed to build the phylogenetic tree using mashtree with the minimum abundance finder.

**Supplemental Figure 4.**
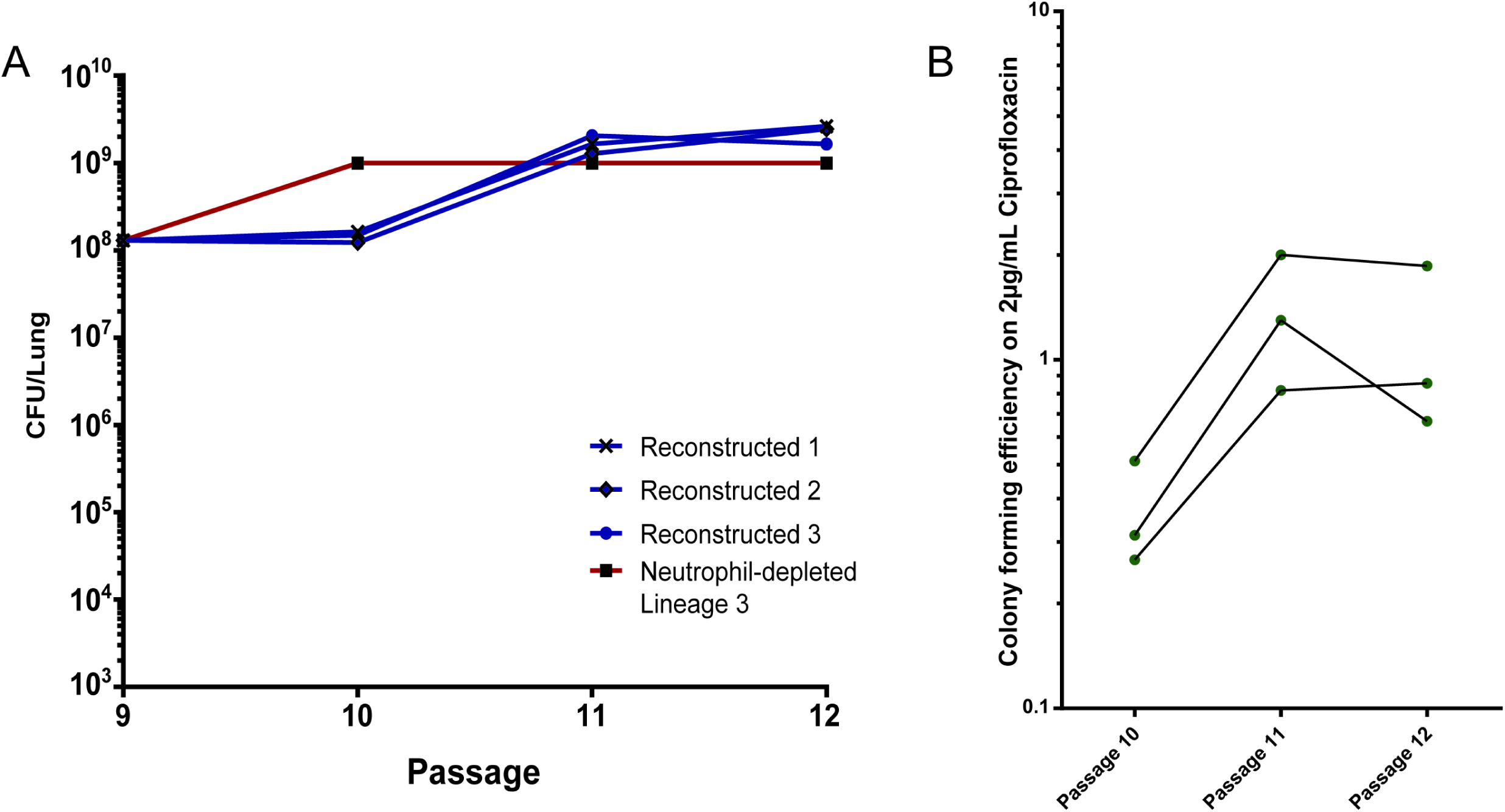
Bacterial load and resistance levels to CIP after re-passaging. A) The bacterial load harvested from the lungs after each passage is plotted against the passage number for each Lineage. B) CIP resistance level of each pool (after each passage) was assayed via quantification of CFU on LB agar supplemented with CIP.

**Supp. Dataset 1.** MLST profiles with 7-gene allelic profiles for 2045 clinical isolates

**Supp. Dataset 2.** All mutations identified in Fig 2 with their detection frequency.

**Supp. Dataset 3.** Multiple alignment of LpxD sequences from 138 clinical isolates.

## Competing Interests Statement

The authors declare that they have no conflict of interest.

## Data availability statement

Sequencing reads that support the findings of this study (Fig 2) are deposited in NCBI SRA with the accession code PRJNA485355. Detailed accession numbers for each sample are listed in Supplementary Table S1. Sequencing reads were analyzed by breseq (https://github.com/barricklab/breseq) and all variants can be found at https://github.com/huoww07/Ab_evolutionary_pathways. Variants were filtered against parent al WT AB17978 and results are attached as Supp. Dataset 2. *A. baumannii* genome data used in this study (Fig 5) are available in the PATRIC database (patricbrc.org) with the sequence IDs listed in Supp. Dataset 1. All other data that support the findings of this study are available on request from the corresponding author RRI.

---

**Table S1.**
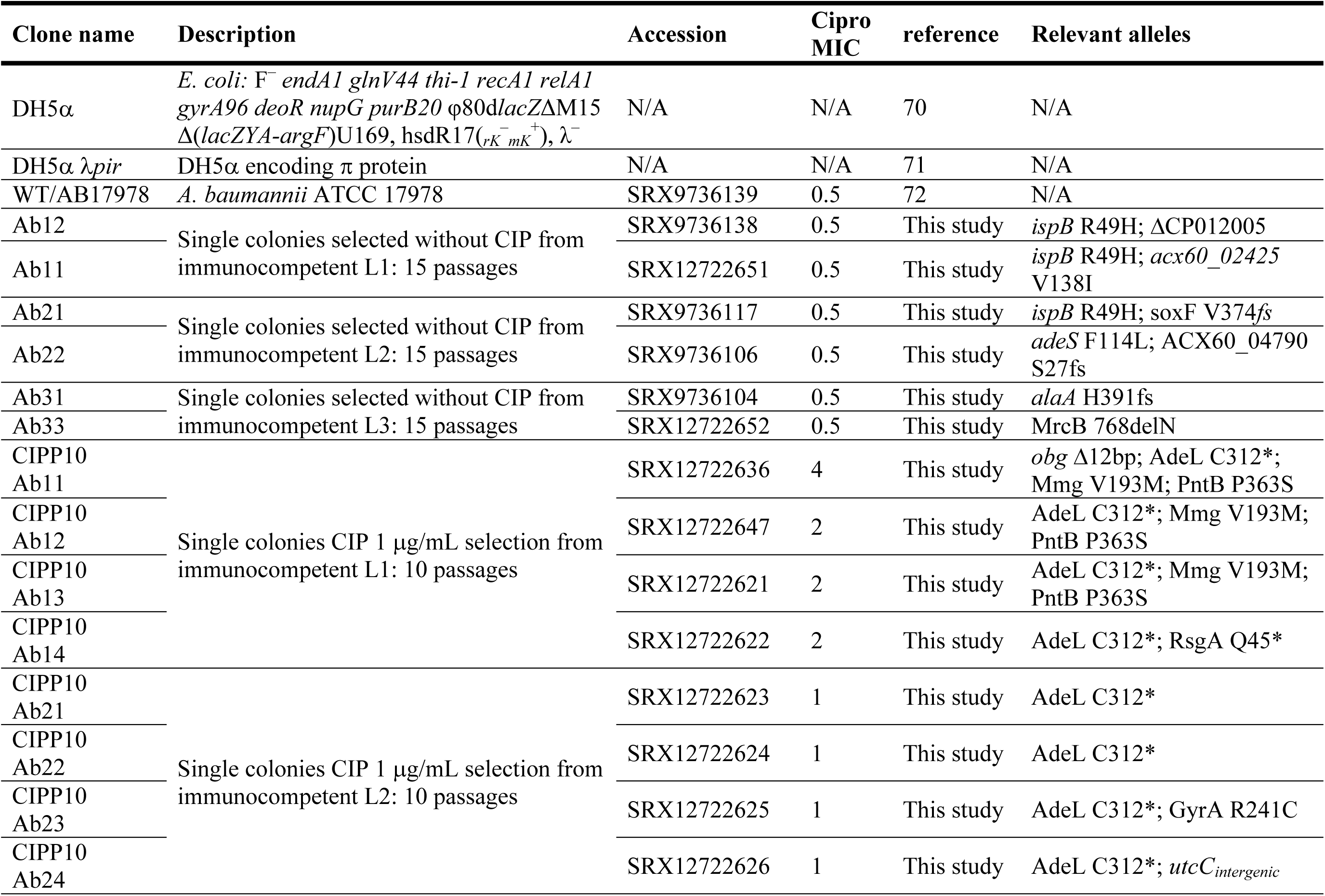
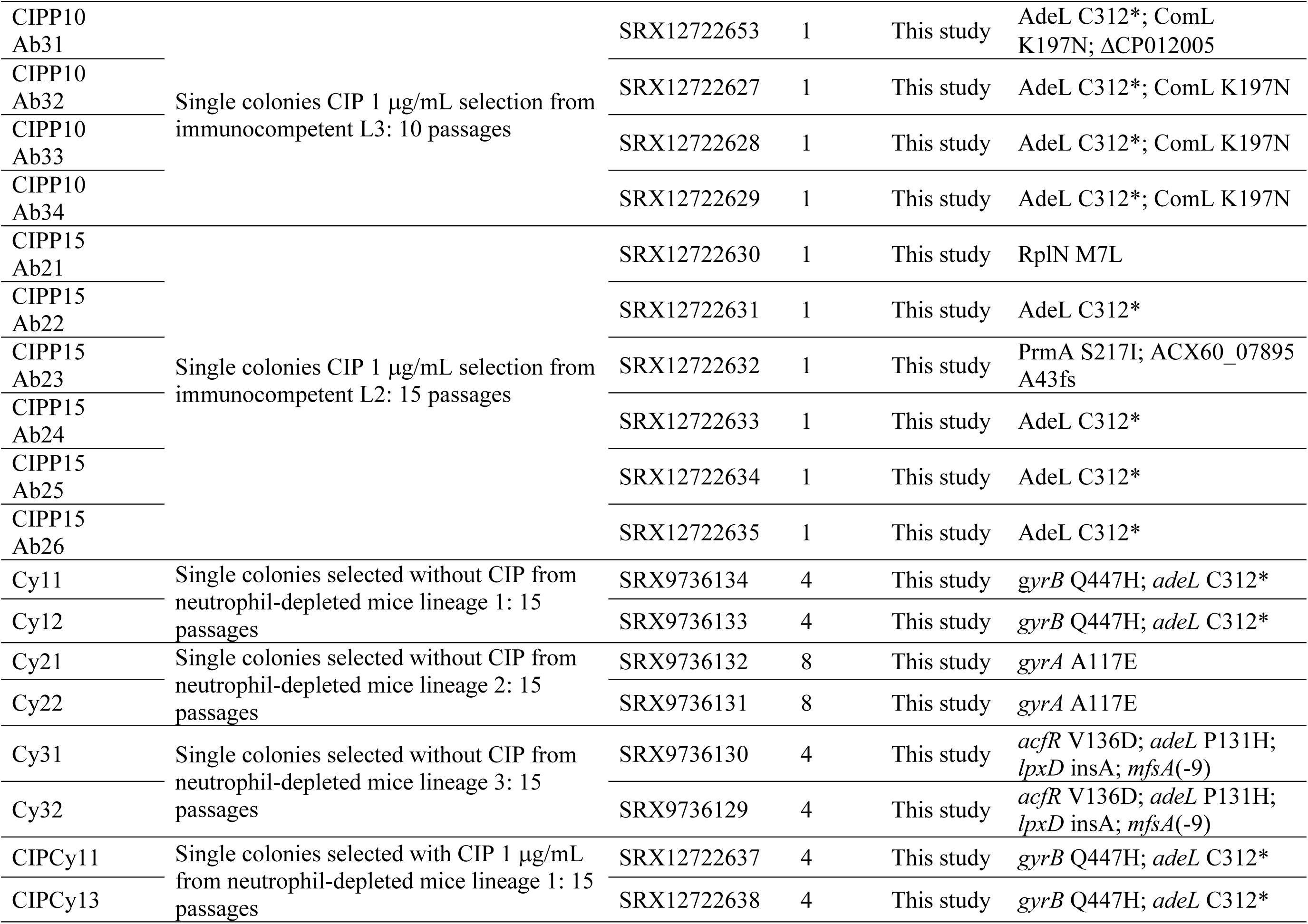
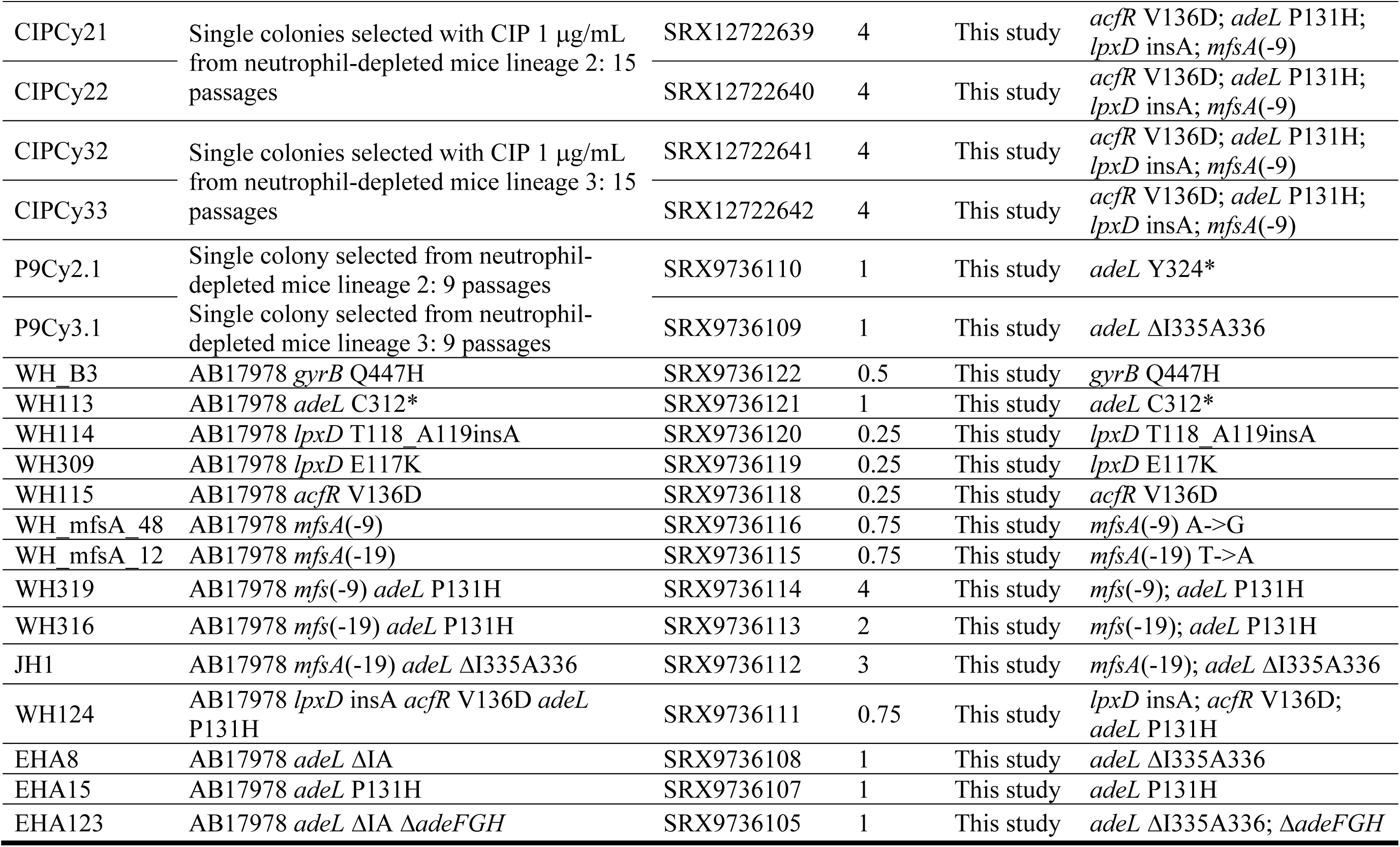
Strains used in this study.

**Table S2.**
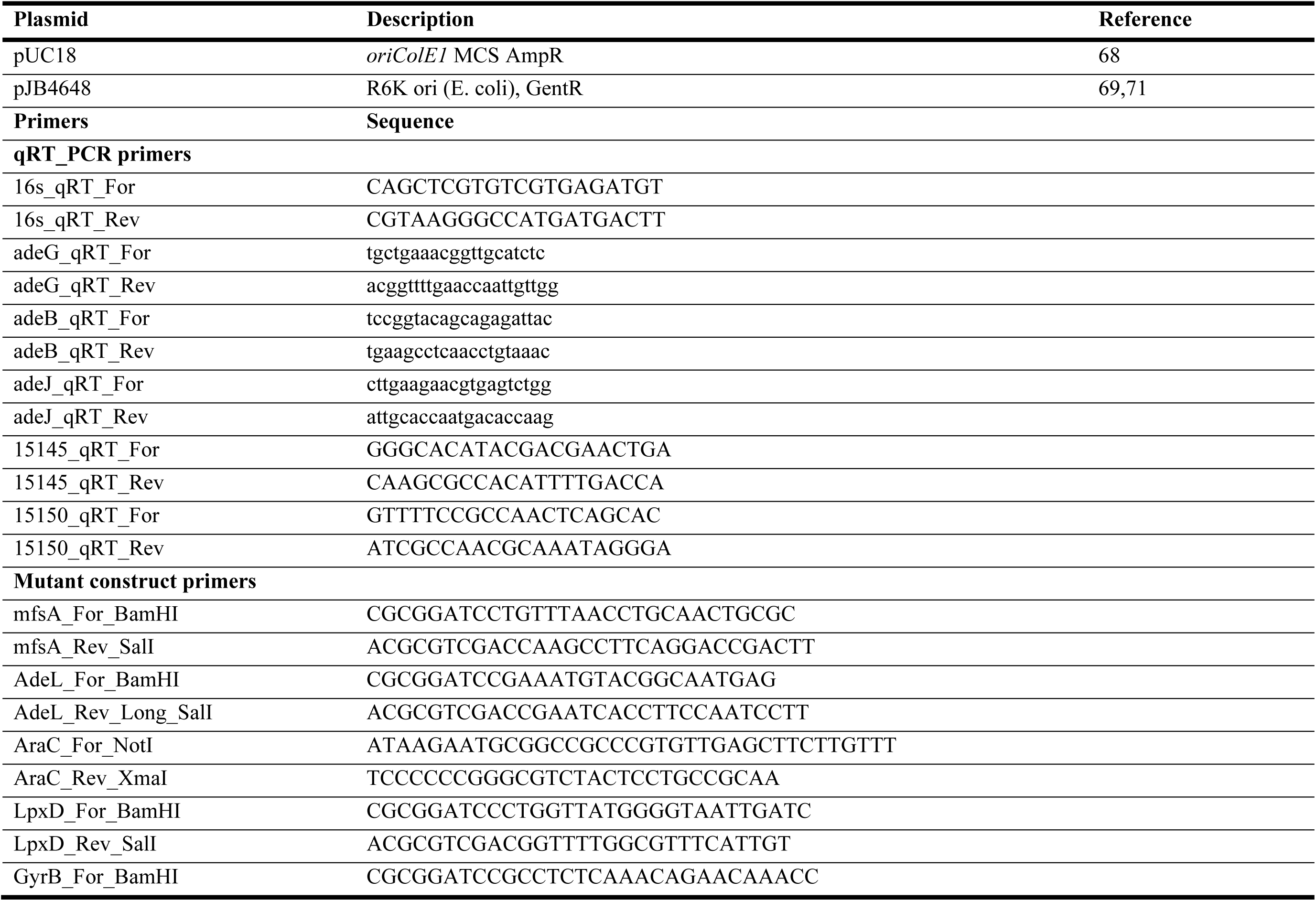
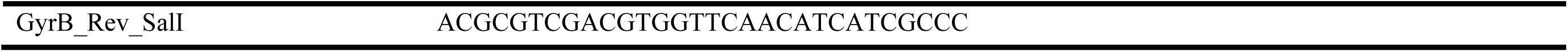
Plasmids and primers used in this study.

**Table S3.** *in vitro* growth rates of LpxD mutants compared to AB17978.

| Time (hr) |  | AB17978 |  |  |  | AB17978::LpxD T118_A119insA |  |  |  | AB17978::LpxD E117K |  |  |  |
| --- | --- | --- | --- | --- | --- | --- | --- | --- | --- | --- | --- | --- | --- |
| Optical<br>density at<br>600nm | 0 | 0.003 | 0.004 | 0.005 | 0.004 | 0.003 | 0.004 | 0.004 | 0.004 | 0.004 | 0.004 | 0.004 | 0.004 |
|  | 1 | 0.015 | 0.015 | 0.015 | 0.013 | 0.009 | 0.011 | 0.011 | 0.011 | 0.014 | 0.014 | 0.014 | 0.013 |
|  | 2 | 0.144 | 0.138 | 0.143 | 0.131 | 0.054 | 0.059 | 0.064 | 0.067 | 0.124 | 0.132 | 0.129 | 0.12 |
|  | 3 | 1.085 | 1.01 | 1.01 | 0.97 | 0.335 | 0.37 | 0.395 | 0.43 | 0.935 | 0.975 | 0.985 | 0.825 |
|  | 4 | 2.47 | 2.68 | 2.445 | 2.38 | 1.16 | 1.27 | 1.355 | 1.39 | 2.36 | 2.475 | 2.475 | 2.27 |
|  | 5 | 3.79 | 3.82 | 3.78 | 3.8 | 2.81 | 2.98 | 2.92 | 3.12 | 3.64 | 3.79 | 3.64 | 3.76 |
|  | 6 | 4.26 | 4.23 | 4.3 | 4.29 | 3.56 | 3.52 | 3.57 | 3.81 | 4.41 | 4.31 | 4.33 | 4.12 |
|  | 7 | 4.5 | 4.59 | 4.67 | 4.64 | 3.95 | 3.97 | 3.99 | 4.09 | 4.69 | 4.4 | 4.47 | 4.55 |
|  | 8 | 4.3 | 4.29 | 4.58 | 4.64 | 4.34 | 4.36 | 4.23 | 4.28 | 4.44 | 4.44 | 4.49 | 4.32 |
| **Doubling time(min)** |  | **17.325** | **17.479** | **17.667** | **17.258** | **21.579** | **21.904** | **21.563** | **21.165** | **17.927** | **17.511** | **17.546** | **18.005** |
| **R<sup>2</sup>** |  | **0.999** | **1** | **0.995** | **0.996** | **1** | **1** | **1** | **0.999** | **0.999** | **0.999** | **0.999** | **0.998** |

**Table S4.** *in vitro* growth rates of the isolates acquired from the *in vivo* passaging experiments.

| Time (hr) |  | AB17978 |  |  | AB17978::*adeL* ΔIA |  |  | Cy11 |  |  | Cy21 |  |  | Cy31 |  |  |
| --- | --- | --- | --- | --- | --- | --- | --- | --- | --- | --- | --- | --- | --- | --- | --- | --- |
| Optical<br>density<br>at<br>600nm | 0 | 0.001 | 0.001 | 0.001 | 0.001 | 0.001 | 0.001 | 0.001 | 0.001 | 0.001 | 0.001 | 0.001 | 0.001 | 0.001 | 0.001 | 0.001 |
|  | 1 | 0.004 | 0.002 | 0.004 | 0.003 | 0.003 | 0.003 | 0.003 | 0.003 | 0.003 | 0.003 | 0.003 | 0.003 | 0.003 | 0.004 | 0.003 |
|  | 2 | 0.02 | 0.021 | 0.025 | 0.013 | 0.015 | 0.014 | 0.012 | 0.011 | 0.011 | 0.014 | 0.012 | 0.013 | 0.017 | 0.015 | 0.015 |
|  | 3 | 0.106 | 0.182 | 0.201 | 0.074 | 0.077 | 0.055 | 0.034 | 0.042 | 0.03 | 0.059 | 0.042 | 0.049 | 0.109 | 0.052 | 0.094 |
|  | 4 | 0.566 | 0.712 | 0.747 | 0.417 | 0.416 | 0.284 | 0.111 | 0.235 | 0.084 | 0.33 | 0.22 | 0.222 | 0.532 | 0.272 | 0.488 |
|  | 5 | 1.6 | 1.65 | 1.8 | 1.22 | 1.2 | 0.99 | 0.65 | 0.9 | 0.51 | 1.02 | 0.93 | 1.95 | 1.23 | 1.03 | 1.24 |
|  | 6 | 2.03 | 2.04 | 2.12 | 1.44 | 1.38 | 1.3 | 1.06 | 1.23 | 0.94 | 1.35 | 1.23 | 1.22 | 1.55 | 1.36 | 1.55 |
|  | 7 | 2.3 | 2.29 | 2.5 | 1.69 | 1.76 | 1.63 | 1.38 | 1.54 | 1.1 | 1.57 | 1.5 | 1.39 | 1.88 | 1.74 | 1.78 |
|  | 8 | 2.56 | 2.55 | 2.7 | 1.85 | 1.83 | 1.89 | 1.51 | 1.73 | 1.37 | 1.78 | 1.61 | 1.57 | 2.01 | 1.91 | 2 |
| Doubling<br>time(min) |  | 21.463 | 17.147 | 19.693 | 20.828 | 21.57 | 23.843 | 27.629 | 23.331 | 28.985 | 22.828 | 23.691 | 21.038 | 20.606 | 24.404 | 20.682 |
| R<sup>2</sup> |  | 0.993 | 0.986 | 0.999 | 0.996 | 0.997 | 0.996 | 0.987 | 0.992 | 0.984 | 0.995 | 0.99 | 0.975 | 0.999 | 0.984 | 0.998 |

## Notes

### Competing Interest Statement

The authors have declared no competing interest.

## References

1. Boucher, H. W. et al. Bad Bugs, No Drugs: No ESKAPE! An Update from the Infectious Diseases Society of America. Clinical Infectious Diseases 48, 1–12, doi:10.1086/595011 (2009).

2. Valentine, S. C. et al. Phenotypic and Molecular Characterization of *Acinetobacter baumannii* Clinical Isolates from Nosocomial Outbreaks in Los Angeles County, California. Journal of Clinical Microbiology 46, 2499–2507, doi:10.1128/jcm.00367-08 (2008).

3. Sievert, D. M. et al. Antimicrobial-Resistant Pathogens Associated with Healthcare-Associated Infections Summary of Data Reported to the National Healthcare Safety Network at the Centers for Disease Control and Prevention, 2009–2010. Infection Control & Hospital Epidemiology 34, 1–14, doi:10.1086/668770 (2015).

4. Karlowsky, J. A., Hoban, D. J., DeCorby, M. R., Laing, N. M. & Zhanel, G. G. Fluoroquinolone-Resistant Urinary Isolates of *Escherichia coli* from Outpatients Are Frequently Multidrug Resistant: Results from the North American Urinary Tract Infection Collaborative Alliance-Quinolone Resistance Study. Antimicrobial Agents and Chemotherapy 50, 2251–2254, doi:10.1128/aac.00123-06 (2006).

5. Azim, A. et al. Epidemiology of bacterial colonization at intensive care unit admission with emphasis on extended-spectrum β-lactamase- and metallo-β-lactamase-producing Gram-negative bacteria – an Indian experience. Journal of Medical Microbiology 59, 955–960, doi:10.1099/jmm.0.018085-0 (2010).

6. Montefour, K. et al. *Acinetobacter baumannii:* an emerging multidrug-resistant pathogen in critical care. Crit Care Nurse 28, 15–25; quiz 26 (2008).

7. Zeighami, H., Valadkhani, F., Shapouri, R., Samadi, E. & Haghi, F. Virulence characteristics of multidrug resistant biofilm forming *Acinetobacter baumannii* isolated from intensive care unit patients. BMC Infectious Diseases 19, doi:10.1186/s12879-019-4272-0 (2019).

8. Dijkshoorn, L., Nemec, A. & Seifert, H. An increasing threat in hospitals: multidrug-resistant *Acinetobacter baumannii*. Nature Reviews Microbiology 5, 939–951, doi:10.1038/nrmicro1789 (2007).

9. Ryan, K. et al. The impact of initial antibiotic treatment failure: Real-world insights in healthcare-associated or nosocomial pneumonia. Journal of Infection 77, 9–17, doi:10.1016/j.jinf.2018.04.002 (2018).

10. Wong, D. et al. Clinical and Pathophysiological Overview of *Acinetobacter* Infections: a Century of Challenges. Clin Microbiol Rev 30, 409–447, doi:10.1128/CMR.00058-16 (2017).

11. Fernandez-Cuenca, F. Relationship between beta-lactamase production, outer membrane protein and penicillin-binding protein profiles on the activity of carbapenems against clinical isolates of *Acinetobacter baumannii*. Journal of Antimicrobial Chemotherapy 51, 565–574, doi:10.1093/jac/dkg097 (2003).

12. Lolans, K., Rice, T. W., Munoz-Price, L. S. & Quinn, J. P. Multicity outbreak of carbapenem-resistant *Acinetobacter baumannii* isolates producing the carbapenemase OXA-40. Antimicrob Agents Chemother 50, 2941–2945, doi:10.1128/AAC.00116-06 (2006).

13. Geisinger, E. & Isberg, R. R. Antibiotic modulation of capsular exopolysaccharide and virulence in *Acinetobacter baumannii*. PLoS Pathog 11, e1004691, doi:10.1371/journal.ppat.1004691 (2015).

14. Aldred, K. J., Kerns, R. J. & Osheroff, N. Mechanism of quinolone action and resistance. Biochemistry 53, 1565–1574, doi:10.1021/bi5000564 (2014).

15. Redgrave, L. S., Sutton, S. B., Webber, M. A. & Piddock, L. J. V. Fluoroquinolone resistance: mechanisms, impact on bacteria, and role in evolutionary success. Trends in Microbiology 22, 438–445, doi:10.1016/j.tim.2014.04.007 (2014).

16. Hujer, K. M. et al. Rapid Determination of Quinolone Resistance in *Acinetobacter spp*. Journal of Clinical Microbiology 47, 1436–1442, doi:10.1128/jcm.02380-08 (2009).

17. Munoz-Price, L. S. & Weinstein, R. A. Acinetobacter infection. N Engl J Med 358, 1271–1281, doi:10.1056/NEJMra070741 (2008).

18. Ankomah, P. & Levin, B. R. Exploring the collaboration between antibiotics and the immune response in the treatment of acute, self-limiting infections. Proc Natl Acad Sci U S A 111, 8331–8338, doi:10.1073/pnas.1400352111 (2014).

19. Wheatley, R. et al. Rapid evolution and host immunity drive the rise and fall of carbapenem resistance during an acute *Pseudomonas aeruginosa* infection. Nat Commun 12, 2460, doi:10.1038/s41467-021-22814-9 (2021).

20. Saroj, S. D., Clemmer, K. M., Bonomo, R. A. & Rather, P. N. Novel Mechanism for Fluoroquinolone Resistance in *Acinetobacter baumannii*. Antimicrobial Agents and Chemotherapy 56, 4955–4957, doi:10.1128/aac.00739-12 (2012).

21. Osińska, A., Harnisz, M. & Korzeniewska, E. Prevalence of plasmid-mediated multidrug resistance determinants in fluoroquinolone-resistant bacteria isolated from sewage and surface water. Environmental Science and Pollution Research 23, 10818–10831, doi:10.1007/s11356-016-6221-4 (2016).

22. Elhosseiny, N. M., Elhezawy, N. B. & Attia, A. S. Comparative proteomics analyses of *Acinetobacter baumannii* strains ATCC 17978 and AB5075 reveal the differential role of type II secretion system secretomes in lung colonization and ciprofloxacin resistance. Microbial Pathogenesis 128, 20–27, doi:10.1016/j.micpath.2018.12.039 (2019).

23. Zuluaga, A. F. et al. Neutropenia induced in outbred mice by a simplified low-dose cyclophosphamide regimen: characterization and applicability to diverse experimental models of infectious diseases. BMC Infect Dis 6, 55, doi:10.1186/1471-2334-6-55 (2006).

24. North, R. J. Cyclophosphamide-facilitated adoptive immunotherapy of an established tumor depends on elimination of tumor-induced suppressor T cells. J Exp Med 155, 1063–1074, doi:10.1084/jem.155.4.1063 (1982).

25. Yoon, E. J. et al. Contribution of resistance-nodulation-cell division efflux systems to antibiotic resistance and biofilm formation in *Acinetobacter baumannii*. mBio 6, doi:10.1128/mBio.00309-15 (2015).

26. Coyne, S., Rosenfeld, N., Lambert, T., Courvalin, P. & Perichon, B. Overexpression of resistance-nodulation-cell division pump AdeFGH confers multidrug resistance in *Acinetobacter baumannii*. Antimicrob Agents Chemother 54, 4389–4393, doi:10.1128/AAC.00155-10 (2010).

27. USCAST. The United States Committee on Antimicrobial Susceptibility Testing (USCAST). Breakpoint tables for interpretation of MIC and zone diameter results. Version 6.0. (2020).

28. Brauner, A., Fridman, O., Gefen, O. & Balaban, N. Q. Distinguishing between resistance, tolerance and persistence to antibiotic treatment. Nat Rev Microbiol 14, 320–330, doi:10.1038/nrmicro.2016.34 (2016).

29. Kussell, E., Kishony, R., Balaban, N. Q. & Leibler, S. Bacterial persistence: a model of survival in changing environments. Genetics 169, 1807–1814, doi:10.1534/genetics.104.035352 (2005).

30. Balaban, N. Q. Persistence: mechanisms for triggering and enhancing phenotypic variability. Curr Opin Genet Dev 21, 768–775, doi:10.1016/j.gde.2011.10.001 (2011).

31. Balaban, N. Q. et al. Definitions and guidelines for research on antibiotic persistence. Nat Rev Microbiol 17, 441–448, doi:10.1038/s41579-019-0196-3 (2019).

32. Liu, J., Gefen, O., Ronin, I., Bar-Meir, M. & Balaban, N. Q. Effect of tolerance on the evolution of antibiotic resistance under drug combinations. Science 367, 200–204, doi:10.1126/science.aay3041 (2020).

33. Papkou, A., Hedge, J., Kapel, N., Young, B. & MacLean, R. C. Efflux pump activity potentiates the evolution of antibiotic resistance across *S. aureus* isolates. Nat Commun 11, 3970, doi:10.1038/s41467-020-17735-y (2020).

34. Santos-Lopez, A., Marshall, C. W., Scribner, M. R., Snyder, D. J. & Cooper, V. S. Evolutionary pathways to antibiotic resistance are dependent upon environmental structure and bacterial lifestyle. eLife 8, doi:10.7554/eLife.47612 (2019).

35. Raetz, C. R. H., Reynolds, C. M., Trent, M. S. & Bishop, R. E. Lipid A Modification Systems in Gram-Negative Bacteria. Annual Review of Biochemistry 76, 295–329, doi:10.1146/annurev.biochem.76.010307.145803 (2007).

36. Lathe, R. & Lecocq, J. P. The firA gene, a locus involved in the expression of rifampicin resistance in *Escherichia coli*. Molecular and General Genetics MGG 154, 53–60, doi:10.1007/bf00265576 (1977).

37. Kelly, T. M., Stachula, S. A., Raetz, C. R. & Anderson, M. S. The firA gene of *Escherichia coli* encodes UDP-3-O-(R-3-hydroxymyristoyl)-glucosamine N-acyltransferase. The third step of endotoxin biosynthesis. J Biol Chem 268, 19866–19874 (1993).

38. Buetow, L., Smith, T. K., Dawson, A., Fyffe, S. & Hunter, W. N. Structure and reactivity of LpxD, the N-acyltransferase of lipid A biosynthesis. Proceedings of the National Academy of Sciences 104, 4321–4326, doi:10.1073/pnas.0606356104 (2007).

39. Jolley, K. A., Bray, J. E. & Maiden, M. C. J. Open-access bacterial population genomics: BIGSdb software, the PubMLST.org website and their applications. Wellcome Open Res 3, 124, doi:10.12688/wellcomeopenres.14826.1 (2018).

40. Blount, Z. D., Lenski, R. E. & Losos, J. B. Contingency and determinism in evolution: Replaying life’s tape. Science 362, doi:10.1126/science.aam5979 (2018).

41. Melnyk, A. H., Wong, A. & Kassen, R. The fitness costs of antibiotic resistance mutations. Evolutionary Applications 8, 273–283, doi:10.1111/eva.12196 (2014).

42. Subedi, D., Vijay, A. K., Kohli, G. S., Rice, S. A. & Willcox, M. Comparative genomics of clinical strains of *Pseudomonas aeruginosa* strains isolated from different geographic sites. Scientific Reports 8, doi:10.1038/s41598-018-34020-7 (2018).

43. Lieberman, T. D. et al. Genetic variation of a bacterial pathogen within individuals with cystic fibrosis provides a record of selective pressures. Nature Genetics 46, 82–87, doi:10.1038/ng.2848 (2013).

44. Lieberman, T. D. et al. Parallel bacterial evolution within multiple patients identifies candidate pathogenicity genes. Nature Genetics 43, 1275–1280, doi:10.1038/ng.997 (2011).

45. Choi, B. et al. Persistence and Evolution of SARS-CoV-2 in an Immunocompromised Host. New England Journal of Medicine 383, 2291–2293, doi:10.1056/NEJMc2031364 (2020).

46. Honsa, E. S. et al. RelA Mutant *Enterococcus faecium* with Multiantibiotic Tolerance Arising in an Immunocompromised Host. mBio 8, doi:10.1128/mBio.02124-16 (2017).

47. Ma, C., Yang, X. & Lewis, P. J. Bacterial Transcription as a Target for Antibacterial Drug Development. Microbiology and Molecular Biology Reviews 80, 139–160, doi:10.1128/mmbr.00055-15 (2016).

48. van Hoek, A. H. A. M. et al. Acquired Antibiotic Resistance Genes: An Overview. Frontiers in Microbiology 2, doi:10.3389/fmicb.2011.00203 (2011).

49. Moyed, H. S. & Bertrand, K. P. hipA, a newly recognized gene of *Escherichia coli* K-12 that affects frequency of persistence after inhibition of murein synthesis. J Bacteriol 155, 768–775, doi:10.1128/JB.155.2.768-775.1983 (1983).

50. Schumacher, M. A. et al. HipBA-promoter structures reveal the basis of heritable multidrug tolerance. Nature 524, 59–64, doi:10.1038/nature14662 (2015).

51. Pontes, M. H., Groisman, E. A. & Garsin, D. A. A Physiological Basis for Nonheritable Antibiotic Resistance. mBio 11, doi:10.1128/mBio.00817-20 (2020).

52. Trastoy, R. et al. Mechanisms of Bacterial Tolerance and Persistence in the Gastrointestinal and Respiratory Environments. Clinical Microbiology Reviews 31, doi:10.1128/cmr.00023-18 (2018).

53. Balaban, N. Q., Merrin, J., Chait, R., Kowalik, L. & Leibler, S. Bacterial persistence as a phenotypic switch. Science 305, 1622–1625, doi:10.1126/science.1099390 (2004).

54. Lewis, K. Persister cells, dormancy and infectious disease. Nat Rev Microbiol 5, 48–56, doi:10.1038/nrmicro1557 (2007).

55. Shah, D. et al. Persisters: a distinct physiological state of E. coli. BMC Microbiol 6, 53, doi:10.1186/1471-2180-6-53 (2006).

56. Nandakumar, M., Nathan, C. & Rhee, K. Y. Isocitrate lyase mediates broad antibiotic tolerance in *Mycobacterium tuberculosis*. Nature Communications 5, doi:10.1038/ncomms5306 (2014).

57. Michiels, J. E., Van den Bergh, B., Verstraeten, N. & Michiels, J. Molecular mechanisms and clinical implications of bacterial persistence. Drug Resistance Updates 29, 76–89, doi:10.1016/j.drup.2016.10.002 (2016).

58. Fisher, R. A., Gollan, B. & Helaine, S. Persistent bacterial infections and persister cells. Nature Reviews Microbiology 15, 453–464, doi:10.1038/nrmicro.2017.42 (2017).

59. Van den Bergh, B., Fauvart, M. & Michiels, J. Formation, physiology, ecology, evolution and clinical importance of bacterial persisters. FEMS Microbiology Reviews 41, 219–251, doi:10.1093/femsre/fux001 (2017).

60. Yoon, E. J., Courvalin, P. & Grillot-Courvalin, C. RND-type efflux pumps in multidrug-resistant clinical isolates of *Acinetobacter baumannii*: major role for AdeABC overexpression and AdeRS mutations. Antimicrob Agents Chemother 57, 2989–2995, doi:10.1128/AAC.02556-12 (2013).

61. Pu, Y. et al. Enhanced Efflux Activity Facilitates Drug Tolerance in Dormant Bacterial Cells. Molecular Cell 62, 284–294, doi:10.1016/j.molcel.2016.03.035 (2016).

62. Harms, A., Maisonneuve, E. & Gerdes, K. Mechanisms of bacterial persistence during stress and antibiotic exposure. Science 354, doi:10.1126/science.aaf4268 (2016).

63. Frimodt-Møller, J. & Løbner-Olesen, A. Efflux-Pump Upregulation: From Tolerance to High-level Antibiotic Resistance? Trends in Microbiology 27, 291–293, doi:10.1016/j.tim.2019.01.005 (2019).

64. El Meouche, I. & Dunlop, M. J. Heterogeneity in efflux pump expression predisposes antibiotic-resistant cells to mutation. Science 362, 686–690, doi:10.1126/science.aar7981 (2018).

65. Nolivos, S. et al. Role of AcrAB-TolC multidrug efflux pump in drug-resistance acquisition by plasmid transfer. Science 364, 778–782, doi:10.1126/science.aav6390 (2019).

66. Peeters, P. et al. The impact of initial antibiotic treatment failure: real-world insights in patients with complicated, health care-associated intra-abdominal infection. Infection and Drug Resistance **Volume** 12, 329–343, doi:10.2147/idr.S184116 (2019).

67. Soriano, A., Mensa, J., Meylan, S., Morata, L. & Kuehl, R. When antibiotics fail: a clinical and microbiological perspective on antibiotic tolerance and persistence of *Staphylococcus aureus*. Journal of Antimicrobial Chemotherapy 75, 1071–1086, doi:10.1093/jac/dkz559 (2020).

68. Meylan, S., Andrews, I. W. & Collins, J. J. Targeting Antibiotic Tolerance, Pathogen by Pathogen. Cell 172, 1228–1238, doi:10.1016/j.cell.2018.01.037 (2018).

69. Karve, S. et al. The impact of initial antibiotic treatment failure: Real-world insights in patients with complicated urinary tract infection. Journal of Infection 76, 121–131, doi:10.1016/j.jinf.2017.11.001 (2018).

70. Wang, E. et al. Pathogenesis of Pneumococcal Pneumonia in Cyclophosphamide-Induced Leukopenia in Mice. Infection and Immunity 70, 4226–4238, doi:10.1128/iai.70.8.4226-4238.2002 (2002).

71. Nielsen, T. B., Yan, J., Luna, B. & Spellberg, B. Murine Oropharyngeal Aspiration Model of Ventilator-associated and Hospital-acquired Bacterial Pneumonia. Journal of Visualized Experiments, doi:10.3791/57672 (2018).

72. Travisano, M. & Lenski, R. E. Long-term experimental evolution in *Escherichia coli*. IV. Targets of selection and the specificity of adaptation. Genetics 143, 15–26, doi:10.1093/genetics/143.1.15 (1996).

73. Geisinger, E., Mortman, N. J., Vargas-Cuebas, G., Tai, A. K. & Isberg, R. R. A global regulatory system links virulence and antibiotic resistance to envelope homeostasis in *Acinetobacter baumannii*. PLoS Pathog 14, e1007030, doi:10.1371/journal.ppat.1007030 (2018).

74. Deatherage, D. E. & Barrick, J. E. Identification of mutations in laboratory-evolved microbes from next-generation sequencing data using breseq. Methods Mol Biol 1151, 165–188, doi:10.1007/978-1-4939-0554-6_12 (2014).

75. Wijers, C. D. M. et al. Identification of Two Variants of *Acinetobacter baumannii* Strain ATCC 17978 with Distinct Genotypes and Phenotypes. Infect Immun 89, e0045421, doi:10.1128/IAI.00454-21 (2021).

76. Choi, Y., Sims, G. E., Murphy, S., Miller, J. R. & Chan, A. P. Predicting the functional effect of amino acid substitutions and indels. PLoS One 7, e46688, doi:10.1371/journal.pone.0046688 (2012).

77. Yanisch-Perron, C., Vieira, J. & Messing, J. Improved M13 phage cloning vectors and host strains: nucleotide sequences of the M13mpl8 and pUC19 vectors. Gene 33, 103–119, doi:10.1016/0378-1119(85)90120-9 (1985).

78. Andrews, H. L., Vogel, J. P. & Isberg, R. R. Identification of linked *Legionella pneumophila* genes essential for intracellular growth and evasion of the endocytic pathway. Infect Immun 66, 950–958, doi:10.1128/IAI.66.3.950-958.1998 (1998).

79. Katz, L. S. et al. Mashtree: a rapid comparison of whole genome sequence files. Journal of Open Source Software 4, doi:10.21105/joss.01762 (2019).

80. Davis, J. J. et al. The PATRIC Bioinformatics Resource Center: expanding data and analysis capabilities. Nucleic Acids Research, doi:10.1093/nar/gkz943 (2019).

